# Leptin Activated Hypothalamic BNC2 Neurons Acutely Suppress Food Intake

**DOI:** 10.1101/2024.01.25.577315

**Authors:** Han L. Tan, Luping Yin, Yuqi Tan, Jessica Ivanov, Kaja Plucinska, Anoj Ilanges, Brian R. Herb, Putianqi Wang, Christin Kosse, Paul Cohen, Dayu Lin, Jeffrey M. Friedman

## Abstract

Leptin is an adipose tissue hormone that maintains homeostatic control of adipose tissue mass by regulating the activity of specific neural populations controlling appetite and metabolism^1^. Leptin regulates food intake by inhibiting orexigenic agouti-related protein (AGRP) neurons and activating anorexigenic pro-opiomelanocortin (POMC) neurons^2^. However, while AGRP neurons regulate food intake on a rapid time scale, acute activation of POMC neurons has only a minimal effect^3–5^. This has raised the possibility that there is a heretofore unidentified leptin-regulated neural population that suppresses appetite on a rapid time scale. Here, we report the discovery of a novel population of leptin-target neurons expressing basonuclin 2 (*Bnc2*) that acutely suppress appetite by directly inhibiting AGRP neurons. Opposite to the effect of AGRP activation, BNC2 neuronal activation elicited a place preference indicative of positive valence in hungry but not fed mice. The activity of BNC2 neurons is finely tuned by leptin, sensory food cues, and nutritional status. Finally, deleting leptin receptors in BNC2 neurons caused marked hyperphagia and obesity, similar to that observed in a leptin receptor knockout in AGRP neurons. These data indicate that BNC2-expressing neurons are a key component of the neural circuit that maintains energy balance, thus filling an important gap in our understanding of the regulation of food intake and leptin action.

## Main

Leptin is an adipose tissue-derived hormone that functions as the afferent signal in a negative feedback loop that maintains homeostatic control of adipose tissue mass^1,6^. Leptin regulates food intake and maintains energy balance in part by inhibiting orexigenic agouti-related protein (AGRP)/neuropeptide Y (NPY) neurons, expressing leptin receptor (LepR), and activating anorexigenic pro-opiomelanocortin (POMC) neurons, also expressing LepR^1,2^. These neurons are located in the arcuate nucleus (ARC) of the hypothalamus, adjacent to the median eminence, the site of a circumventricular organ that has fenestrated capillaries enabling signaling without transit across the blood-brain barrier. POMC is a protein precursor that is proteolytically cleaved to generate alpha-melanocyte stimulating hormone (aMSH) which reduces food intake by activating the melanocortin 4 receptor (MC4R) which is widely distributed throughout the brain^7,8^. AGRP/NPY neurons increase food intake via projections to many of the same sites as POMC neurons where the AGRP protein diminishes aMSH signaling at the MC4 receptor^7,8^. AGRP/NPY neurons also directly inhibit POMC neurons^9^. These findings have suggested that food intake is reciprocally regulated by these two neural populations. This proposed ‘yin-yang’ dynamic between these neural populations is a consistent feature in nearly all models of how the homeostatic control of food intake and body weight is maintained^10^. However, while leptin acutely reduces food intake, particularly in ob/ob mice, this response cannot be fully accounted for by modulation of AGRP/NPY or POMC neurons^11,12^. Moreover, the functional effects and dynamics of POMC and AGRP/NPY neurons diverge in several important respects and several lines of evidence have suggested that there may be additional leptin-responsive populations that are crucial for leptin-driven control of food intake and body weight^13,14^.

First, while activating AGRP/NPY neurons rapidly leads to food seeking and consumption, activating POMC neurons has only a minimal effect to acutely reduce food intake^3–5^. Second, mutations in POMC, POMC processing enzymes, or the MC4R cause obesity^7,8,15^, whereas mutations in AGRP and NPY have not been reported to alter body weight although ablation of these neurons has been reported to cause anorexia^16–20^. In addition, while activation of AGRP/NPY neurons is associated with negative valence, POMC neuron activation is not associated with positive valence but rather also elicits negative valence^21,22^. The role of these populations in leptin signaling has also been evaluated and while a knockout of the LepR in AGRP/NPY neurons of adult mice causes extreme obesity, deletion of the LepR from adult POMC neurons has only a minimal effect^23^. Finally, acute activation of AGRP/NPY neurons alters glucose metabolism, a response not observed with acute manipulation of POMC neurons^24^. Thus, while POMC and AGRP/NPY neurons have opposite effects on weight, their effects are not equivalent in other respects, indicating that they are not precise counterparts. This has raised the possibility that there might be a missing population of LepR-expressing neurons that acutely suppresses food intake with a time course similar to the effects of AGRP/NPY neurons.

To address this, we conducted a systematic profiling of the transcriptomes of ARC neurons using single nucleus RNA sequencing (snRNA-seq) and identified a novel population of LepR-expressing ARC neurons that co-express the basonuclin 2 (*Bnc2*) gene. Further studies revealed that BNC2 neurons rapidly induce satiety with similar kinetics to the activation of feeding after AGRP/NPY neural activation. This discovery adds an important new cellular component to the neural network that regulates hunger and satiety and sheds new light on how leptin regulates energy balance.

## Results

### Identification of novel LepR-expressing neurons in the ARC

We performed snRNA-seq on ARC that was micro-dissected from adult mice (**Fig. 1a**). Nuclei were isolated, stained with a fluorescent antibody to NeuN, a specific marker of neuronal nuclei, and subjected to fluorescent activated sorting followed by single nucleus sequencing using the 10X single-cell analysis platform^25^. A total of 3,557 cells were profiled, of which 3,481 were identified as neurons based on the expression levels of neuronal markers including *Tubb3*, *Snap25*, *Syt1*, and *Elavl4,* and the absence of non-neuronal markers (**Extended Data Fig. 1a,b**). Leiden clustering analysis identified 21 distinct neuron clusters (**Fig. 1b,c**). To confirm that these neurons were from the ARC rather than from adjacent regions, we analyzed the clusters for specific marker genes that define known ARC populations. This validation process confirmed that 19 of the clusters were indeed located within the ARC with the exception of clusters 9 and 15, which express *Nr5a1*, indicating their origin from the adjacent ventromedial hypothalamus (VMH) region (**Fig. 1c**)^26,27^. These clusters included one AGRP/NPY (Cluster 0), two POMC-expressing (Clusters 3 and 12) neurons, and 16 additional clusters (**Fig. 1d**). Clusters 0 and 3 expressed the LepR as did Cluster 17 which did not express AGRP, NPY, or POMC but instead was characterized by the expression of the *Bnc2* gene (**Fig. 1d**). Several of the genes in BNC2 Cluster 17 including *Lepr* were also found in a single cluster (n11.Trh/Cxcl12) in a previously published dataset of ARC neuron types^26^ (**Extended Data Fig. 1c**). However, *Bnc2* was not identified in this cluster and thyrotropinreleasing hormone (*Trh)*, the canonical marker, was also expressed in other clusters that did not express LepR^26^. Thus, BNC2 serves as a specific marker for a novel LepR cluster that does not include AGRP, NPY or POMC.

**Fig. 1:**
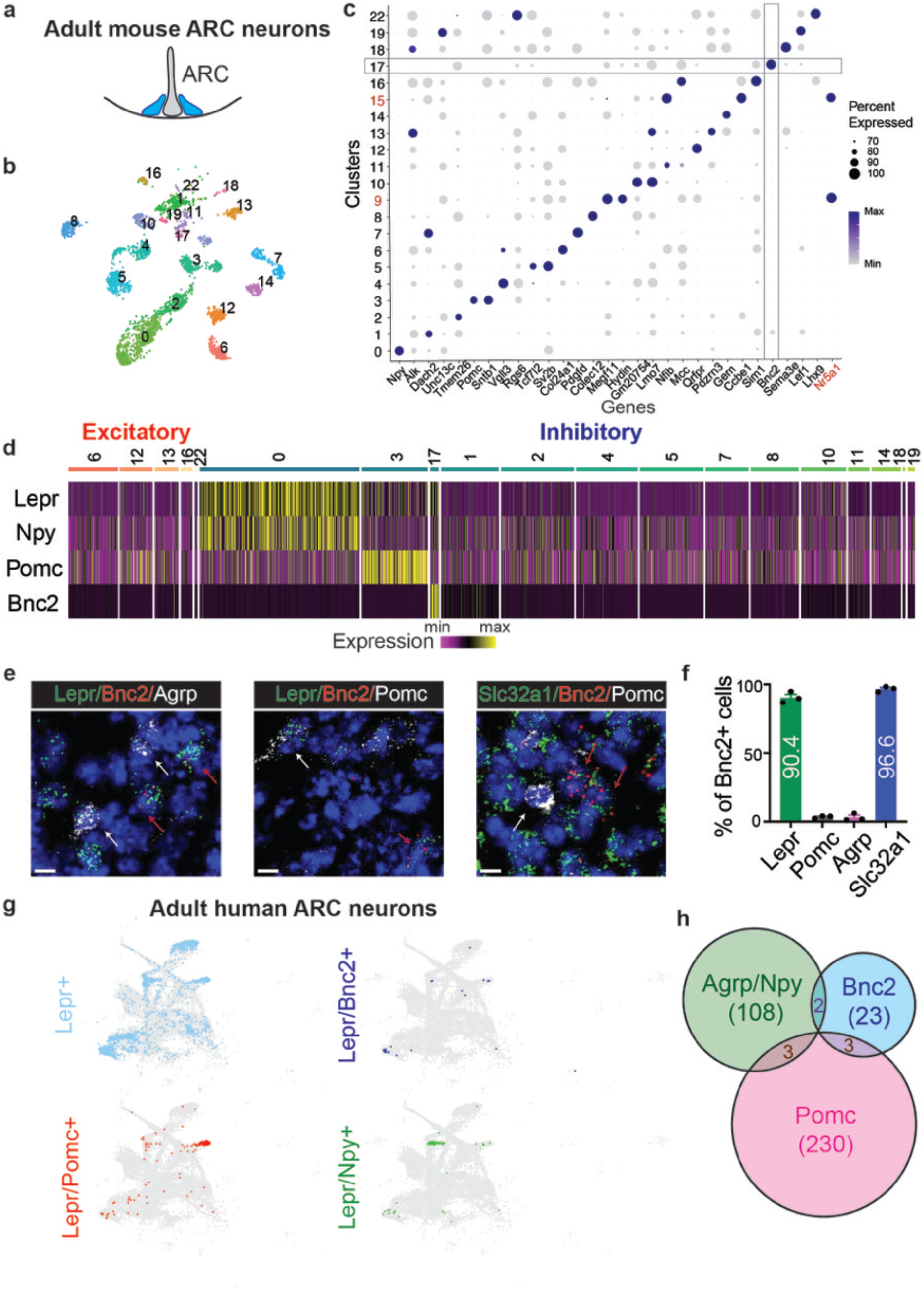
Identification of novel LepR-expressing neurons in the ARC. **a**, Micro-dissection of mouse ARC. **b,** A uniform manifold approximation and projection (UMAP) plot of neuronal nuclei from the ARC (N = 6 adult male WT mice). **c,** Dot plot of maker genes of individual neuronal clusters and *Nr5a1* gene. **d,** Heatmap to plot the expression of indicated genes across individual clusters. Clusters were categorized into excitatory (*Slc17a6* positive) and inhibitory (*Slc321a*/*Gad1*/*Gad2* positive) neurons. **e,** RNA ISH against *Lepr*, *Bnc2*, *Agrp*, *Pomc*, *Slc32a1* in ARC of adult male WT mice. White arrow: *Agrp or Pomc positive* cells; Red arrow: *Bnc2 positive* cells. Scale bar, 10 µm. **f,** Percentage of *Bnc2*-positive cells co-expressing *Lepr*, *Pomc*, *Agrp*, and *Slc32a1* (N = 3 mice). **g,** UMAP plots of *Lepr*/*Pomc*, *Lepr*/*Bnc2*, and *Lepr*/*Npy* neurons in adult human ARC neurons. **h,** Colocalization of *Lepr*/*Bnc2*, *Lepr*/*Agrp*/*Npy*, and *Lepr*/*Pomc* neurons in adult human ARC neurons. Data are presented as mean ± SEM.

The total number of cells in the BNC2 cluster (Cluster 17, n=35) was lower than those in the AGRP cluster (Clusters 0, n=650) or the POMC cluster (Cluster 3, n=271). However, the level of LepR expression in the cells in this cluster was significantly higher than in the POMC cluster and equivalent to that in the AGRP cluster (Cluster 0: 1.32; Cluster 3: 0.52; Cluster 17: 1.09). We confirmed the co-expression of LepR and BNC2 using multiplex RNA in situ hybridization of adult mouse hypothalamus. The results showed that over 90% of *Bnc2* cells in the ARC co-expressed *Lepr* (90.4%) and *Slc32a1* (96.6%), a marker for inhibitory GABAergic neurons while less than 5% co-expressed either *Agrp* (2.6%) and *Pomc* (3.3%) (**Fig. 1e,f**). We also determined whether BNC2 co-localized with LepR in human hypothalamus ARC and found a significant overlap between *Bnc2* and *Lepr* (Fisher’s exact test, p=0.0218, **Fig. 1g**). Here again, the LepR/BNC2 cells in human hypothalamus were distinct from AGRP/NPY and POMC neurons, as 82.1% (23 out of 28) of LepR/BNC2 neurons were not co-localized with *Agrp*, *Npy*, or *Pomc* (**Fig. 1h**). Altogether, these results show that BNC2 neurons are a novel population of LepR-expressing neurons in mouse and human that are non-overlapping with the previously characterized LepR AGRP/NPY and POMC populations. Spatial transcriptome data from the Allen Brain Institute confirms hypothalamic expression of BNC2 with limited expression elsewhere in the brain other than in the cortex and hippocampus^28^.

### Activation of BNC2 neurons by leptin, refeeding, and food cues

In order to further study the dynamics and function of BNC2 neurons in ARC, we generated a BNC2-P2A-iCre knockin mouse line (referred to hereafter as BNC2-Cre) by inserting a P2a-iCre fusion in frame at the C-terminus of the BNC2 protein^29^ (**Extended Data Fig. 2a,b**). The correct insertion site of iCre was confirmed using PCR of genomic DNA (**Extended Data Fig. 2c**). Eutopic expression of Cre in BNC2 neurons in ARC was verified by injecting an AAV virus with a Cre-dependent mCherry into the ARC of adult BNC2-Cre mice. This showed nearly complete co-localization of mCherry and *Bnc2* RNA (**Extended Data Fig. 2d**). Leptin binding to LepR activates Janus tyrosine kinase 2 (Jak2), leading to STAT3 phosphorylation, a canonical marker of leptin activation^30^ and we confirmed leptin signaling in BNC2 neurons by immunostaining for pSTAT3. An AAV expressing a DIO-mCherry cassette was injected into the ARC of adult BNC2-Cre mice and animals subsequently received a single injection of leptin (3 mg/kg) after an overnight fast. This resulted in a significant increase in pSTAT3 in mCherry labeled BNC2 neurons at 3 hours, not seen in control animals that received PBS injections (**Fig. 2a,b**). Refeeding mice after an overnight fast also led to a significant increase of pSTAT3 levels in BNC2 neurons (**Fig. 2a,b**). Consistent with this we also found that the levels of cFos, a neuron activity marker ^31^, significantly increased in BNC2 neurons following refeeding or leptin injection (**Fig. 2c,d**).

**Fig. 2:**
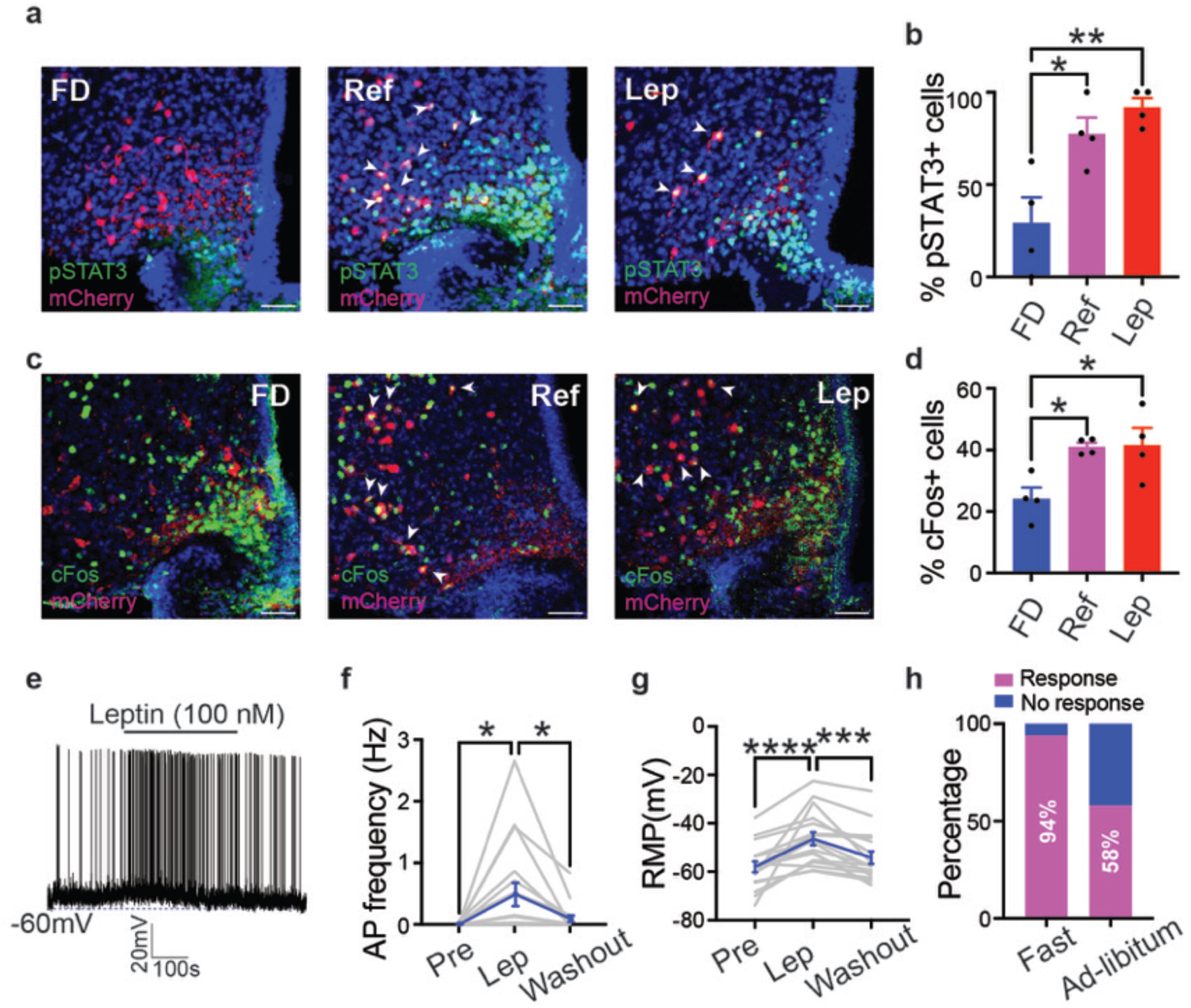
Activation of BNC2 neurons by leptin and feeding. **a,c,** Brain slices from adult male BNC2-Cre mice, injected with DIO-mCherry into ARC, were subjected to pSTAT3 **(a)** and cFos **(c)** immunostaining under three conditions: overnight fasting (FD), overnight fasting followed by refeeding with chow for 3 hours (Ref), and overnight fasting followed by leptin injection (3mg/kg) for 3 hours (Lep). Scale bar, 50µm. **b,d,** Percentage of pSTAT3/mCherry double-positive cells **(b)** or cFos/mCherry double-positive cells **(d)** in mCherry positive cells (N = 4 mice per group). **e,** Spontaneous APs of BNC2 neurons in ARC from overnight fasted adult male BNC2-Cre mice (injected with DIO-GFP). Leptin (100 nM) was added to the bath solution at the indicated time window. **f,g,** Spontaneous AP frequency **(f)** and RMP **(g)** of BNC2 neurons before, during, and after leptin application (N = 17 cells from 4 mice per group). **h,** Percentage of leptin-responsive BNC2 neurons under the overnight fasting condition and ad-libitum-fed condition. Data are presented as mean± SEM. *p<0.05; **p<0.01; ***p<0.001; ****p<0.0001. Details of the statistical analysis are provided as Source Data.

We then performed whole-cell patch-clamp recordings in hypothalamic slices prepared from adult BNC2-Cre mice injected with a DIO-EGFP AAV in the ARC. Application of leptin (100 nM) to brain slices prepared from either fasted or ad-libitum-fed mice resulted in a substantial increase in action potential (AP) firing rates in GFP labeled BNC2 neurons (Fast: 0.01±0.0.005 Hz and 0.488±0.188 Hz before and after leptin, respectively, p=0.0157; Fed: 0.014±0.012 Hz and 0.235±0.118 Hz before and after leptin, respectively, p=0.0062) and a significant depolarization (Fast: −58.02±2.24 mV and −46.41±2.69 mV before and after leptin, respectively, p < 0.0001; Fed: −50.13±1.92 mV and −44.92±2.83 mV before and after leptin, respectively, p=0.0074. **Fig. 2e-g, Extended Data Fig.3**). These effects were reversible after leptin washout (**Fig. 2e-g, Extended Data Fig.3**). In addition, a higher percentage of cells responded to leptin in the slices prepared after an overnight fast (16 out of 17) compared to those responding in slices prepared from ad-libitum-fed mice (7 out of 12) (**Fig. 2h**). The greater effect of leptin on neurons from fasted hypothalamus is consistent with the cFos data showing that the baseline activity of BNC2 neurons is lower after an overnight fast (**Fig. 2d**).

To assess the in vivo responses, we employed fiber photometry to record the effect of food cues on the activity of BNC2 neurons. We injected a DIO-GCamp6s AAV into the ARC of adult BNC2-Cre mice and implanted a fiber above the ARC to record neural activity. Mice were fasted overnight after which an inedible plastic tube, a pellet of standard chow, or peanut butter was provided. The inedible tube did not have a discernible effect on BNC2 neuron activity (**Fig. 3a-c, Extended Data Fig. 4a-c**). However, exposure to standard chow robustly activated BNC2 neurons within seconds relative to the plastic tube (Delta F/F: 0.08±0.13 and 2.83±0.99 for tube and chow, respectively; p=0.0078, **Fig. 3a-c, Extended Data Fig. 4a-c**) and exposure to peanut butter, which has a significantly higher fat content, elicited a significantly greater response than did chow (Delta F/F: 4.44±1.36 for peanut butter, p=0.0078, **Fig. 3a-c, Extended Data Fig. 4a-c**). In contrast, BNC2 neurons from ad-libitum-fed mice showed little or no response to the inedible tube and standard chow (**Fig. 3d,e, Extended Data Fig. 4d-f**) while peanut butter still significantly activated BNC2 neurons (Delta F/F: 0.001±0.322 and 2.764±1.093 for chow and peanut butter, respectively; p=0.0078; **Fig. 3d,e, Extended Data Fig. 4d-f**). These results show that the activity of BNC2 neurons is increased by the presence of food and that this is augmented by palatability and sensitive to nutritional state (i.e; fed vs fasted).

**Fig. 3:**
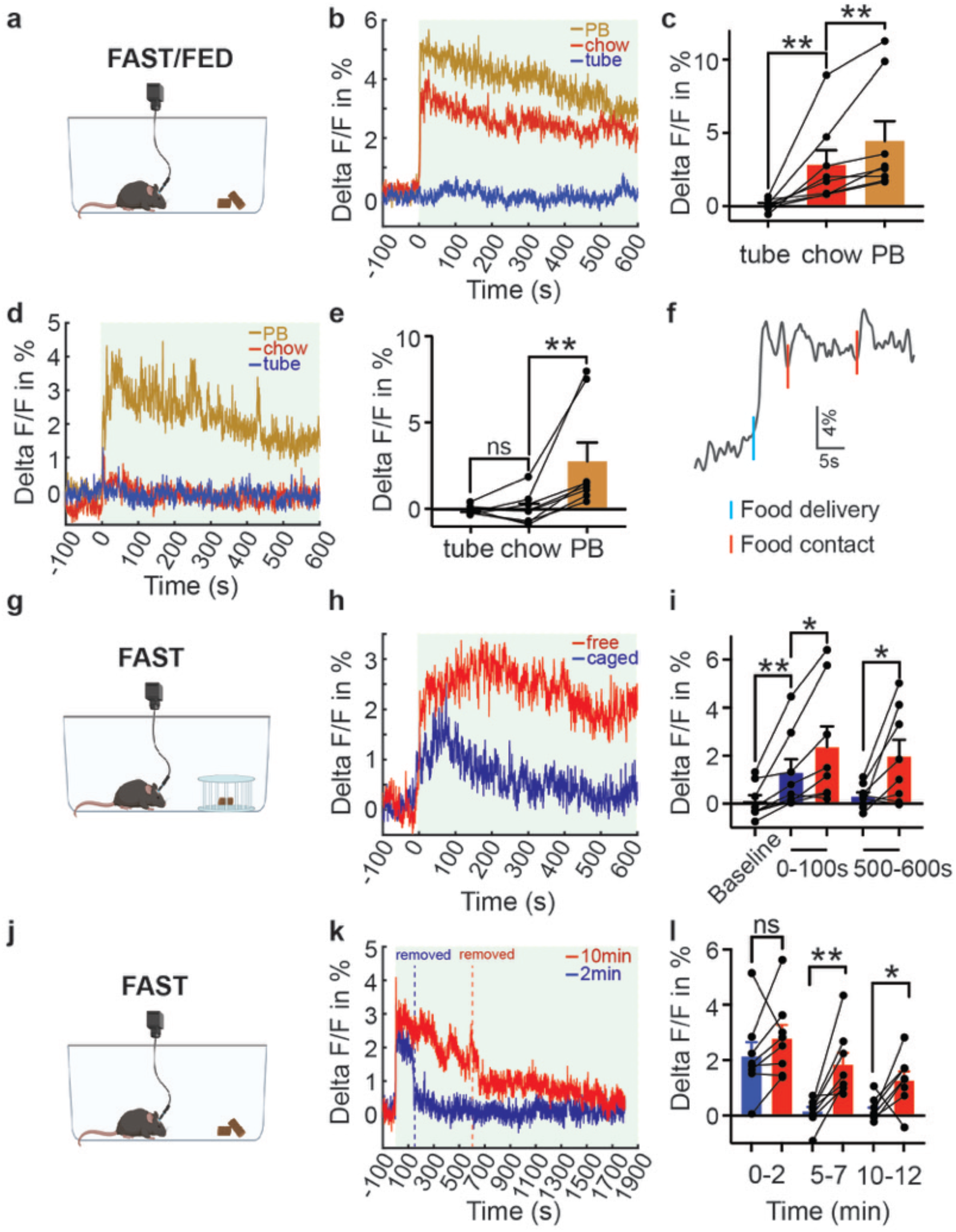
Rapid responses of BNC2 neurons to food cues. **a-f,** Recordings from mice expressing GCaMP6s in BNC2 neurons presented with an inedible plastic tube, chow, or peanut butter (PB). **b,** Average traces of calcium signals in fasted mice aligned to the time of presentation. **c,** Quantification of fluorescence changes within 0-300 second timeframe in b (N = 8 mice per group). **d,** Average traces in ad-libitum-fed mice aligned to the time of presentation. **e,** Quantification of fluorescence changes within 0-300 second timeframe ind (N = 8 mice per group). **f,** Individual trace of one overnight fasted mouse presented with PB. **g-i,** Recordings from overnight fasted mice presented with a chow inside or outside the container. **h,** Average traces aligned to the time of presentation. **i,** Quantification of fluorescence changes within 0-100 second and 500-600 second timeframes in h (N = 8 mice per group). **j-1,** Recordings from overnight fasted mice presented with a chow for 2 minutes or 10 minutes. **k,** Average traces aligned to the time of presentation. I, Quantification of fluorescence changes in the indicated timeframes in **k** (N = 8 mice per group). Data are presented as mean ± SEM. ns, not significant; *p<0.05; **p<0.01. Details of the statistical analysis are provided as Source Data.

We further noted that the increased fluorescence of BNC2 neurons preceded the initiation of food intake suggesting that they were activated by sensory cues (**Fig. 3f, Supplementary Video1**). We tested this directly by presenting mice with standard chow in a container that allowed them to see and smell the pellet without being able to consume it (**Fig. 3g**). Under these conditions, BNC2 neurons still showed significant activation albeit of lesser magnitude and duration compared to the experiments in which the food was not caged (Delta F/F: 1.287±0.567 and 2.354±0.87 for caged and free, respectively; p=0.0156, **Fig. 3h,i, Extended Data 4g,h**). This suggests BNC2 neurons respond to sensory cues and that accessibility to food further modulates the strength and duration of neural activation. Finally, we examined whether the removal of food extinguished the neural activation seen after food presentation. In this experiment, mice that had been fasted overnight were presented with accessible chow which was removed 2 or 10 minutes after it was provided. Removal of food led to a rapid decrease in the activity of BNC2 neurons (**Fig. 3j-l, Extended Data 4i-j**). In contrast, the BNC2 neurons remained active when food was continuously present. (**Fig. 3j-l, Extended Data 4i-j**).

### BNC2 neurons acutely suppress food intake

We next assessed the function of BNC2 neurons using chemogenetics and optogenetics. AAVs expressing a Cre-dependent stimulatory hM3Dq (Gq-coupled human muscarinic M3 designer receptors exclusively activated by designer drugs, DREADD) or an inhibitory hM4Di DREADD were injected into the ARC of BNC2-Cre mice. Mice receiving a virus expressing mCherry and given CNO, as well as mice receiving the DREADD and given PBS were used as controls in these and subsequent experiments. We then selectively activated or inhibited these neurons using clozapine-N-oxide (CNO)^32^. Activating BNC2 neurons at the beginning of the dark cycle resulted in a significant reduction in food intake and body weight compared to control mice (3h food intake: 1.1±0.11 g and 0.52±0.08 g for hM3Dq-PBS and hM3Dq-CNO, respectively;, p=0.0177; 3h body weight change, 0.5±0.04 g and −0.55±0.19 g for hM3Dq-PBS and hM3Dq-CNO, respectively; p<0.0001, **Fig. 4a,b, Extended Data Fig. 5a**). In contrast, silencing BNC2 neurons during the light cycle significantly increased food consumption and body weight (3h food intake: 0.15±0.03 g and 0.51±0.06 g for hM4Di-PBS and hM4Di-CNO, respectively; p=0.0008; 3h body weight change, −0.16±0.08 g and 0.34±0.09 g for hM4Di-PBS and hM3Dq-CNO, respectively; p=0.0002, **Fig. 4c,d**).

**Fig. 4:**
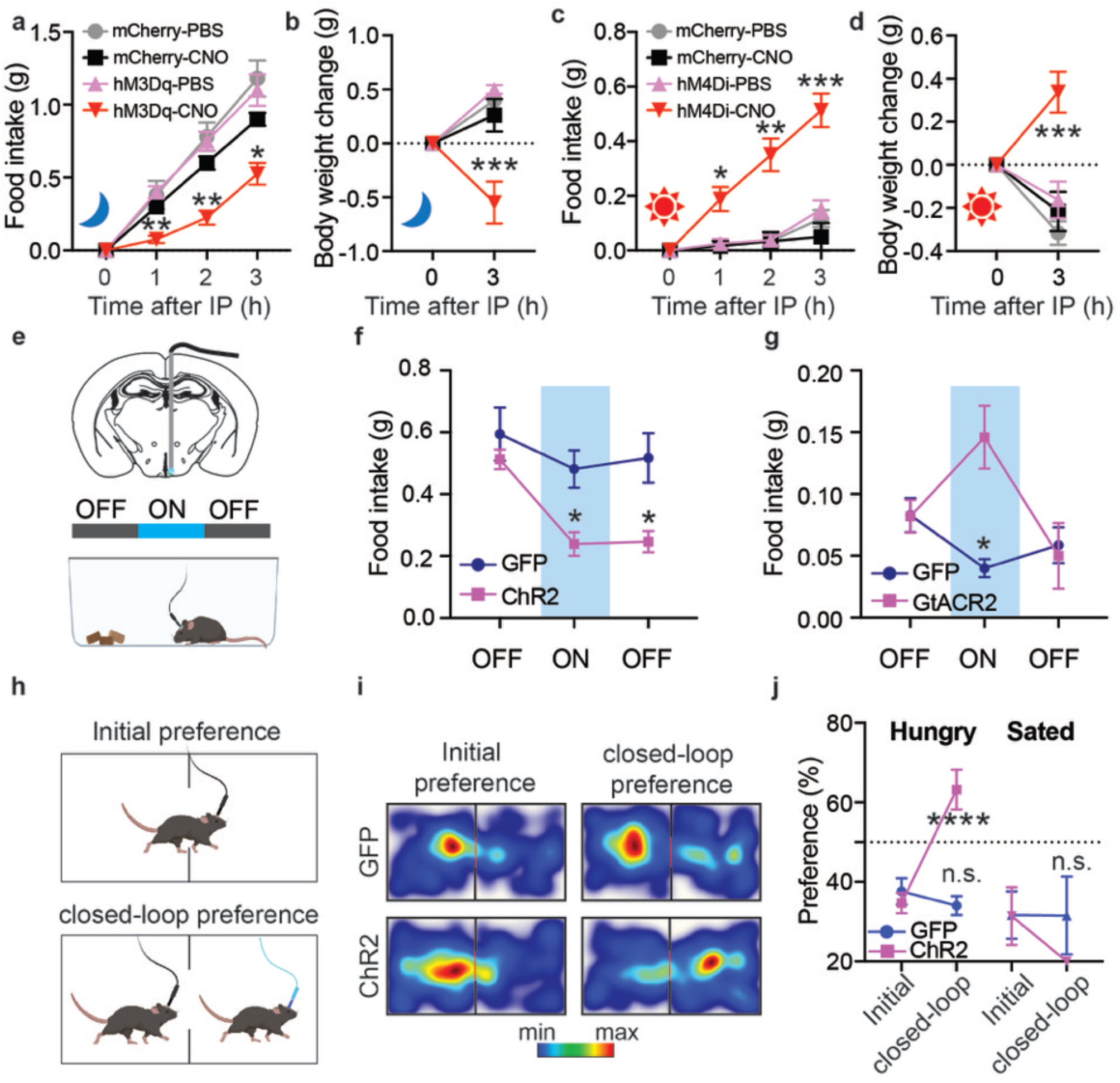
Rapid and sustained satiety driven by BNC2 neurons. **a, b,** Hourly food intake **(a)** and body weight change at 3 hours **(b)** of each group of mice receiving PBS or CNO injection upon entering the night dark phase (N = 5 mCherry-injected mice, N = 4 hM3Dq-injected mice). **c, d,** Hourly food intake (c) and body weight change at 3 hours **(d)** of each group of mice receiving PBS or CNO injection in the daytime (N = 6 mCherry-injected mice, N = 8 hM4Di-injected mice). **e,** Unilateral injection of AAV viruses into the ARC region of adult male BNC2-Cre mice followed by fiber optic implant. **f,** Food intake of mice before, during, and after the activation of laser stimulation in overnight-fasted mice (N = 7 GFP-injected mice, N = 12 ChR2-injected mice). **g,** Food intake of mice before, during, and after the activation of laser stimulation in ad-libitum-fed mice in the daytime (N = 7 GFP-injected mice, N = 5 GtACR2-injected mice). **h-j,** Closed-loop place preference assay to assess the valence of BNC2 neurons. i, Heatmaps of time spent in each chamber of overnight fasted adult male BNC2-Cre mice injected with GFP or ChR2 during the initial phase and the closed-loop paired phase. **j,** Quantification of place preference (time spent in the pair-stimulated chamber, N = 6 GFP-injected mice, N = 8 ChR2-injected mice). ns, not significant; *p<0.05; **p<0.01; ***p<0.001; ****p<0.0001. Details of the statistical analysis are provided as Source Data.

We next evaluated the dynamics of the feeding response using optogenetics^33^. AAVs with Cre-dependent versions of the activating channeldhodopsin-2 (ChR2)-GFP or inhibitory soma-targeted *Guillardia theta* anion-conducting channelrhodopsins (stGtACR2)-FusionRed were injected into the ARC of BNC2-cre mice after which an optical fiber was implanted over the ARC^33,34^. (**Fig. 4e, Extended Data Fig. 5b**). Optogenetic activation of BNC2 neurons at a frequency of 20 Hz for 20 minutes led to a significant reduction in food intake in mice that were fasted overnight compared to controls expressing GFP and the decreased food intake persisted for up to 20 mins even after the cessation of light stimulation (**Fig. 4f**). Activating ARC BNC2 neurons did not affect locomotion (**Extended Data Fig. 5c**). Consistent with the results from chemogenetic inhibition, optogenetic silencing of BNC2 neurons at 20 Hz significantly increased food intake in mice during the light cycle but this did not persist when the photoinhibition ceased (**Fig. 4g**).

We also used optogenetics to assess the effect of BNC2 neural activation on valence using a real-time place preference assay (**Fig. 4h**). We found a significant preference of mice for the chamber associated with BNC2 activation after an overnight fast compared to GFP expressing controls (**Fig. 4i,j**). However, there was no preference for the photostimulation-paired chamber in sated mice that were tested in the light cycle (**Fig. 4j, Extended Data Fig. 5d**). Food deprivation leads to the activation of AGRP/NPY neurons which is associated with negative valence raising the possibility that BNC2 activation is associated with positive valence because these neurons inhibit AGRP/NPY neurons^21,35^. We evaluated this possibility by combining optogenetic activation of BNC2 neurons with electrophysiologic recordings from AGRP/NPY neurons in hypothalamic slices.

### BNC2 neurons monosynaptically inhibit AgRP/NPY neurons

A Flp-dependent mCherry AAV (fDIO-mCherry) and a Cre-dependent ChR2-GFP AAV (DIO-ChR2-GFP) were co-injected into the ARC of *BNC2-Cre*::*NPY-FlpO* mice^36^. We found dense GFP expression in BNC2 terminals within the ARC, in proximity to the mCherry expressing AGRP/NPY soma (**Fig. 5a**). This finding, the positive valence of BNC2 activation after fasting and the observation that BNC2 neurons express Vgat (*Slc32a1*, **Fig. 1e,f**) raised the possibility that BNC2 neurons might directly inhibit AGRP/NPY neurons. We evaluated this using ChR2-assisted circuit mapping (CRACM) on brain slices^37,38^. Evoked inhibitory postsynaptic currents (oIPSCs) were recorded in mCherry-labeled AGRP/NPY neurons before and after optogenetic activation of BNC2 neurons (**Fig. 5b**). We found synchronized oIPSCs after light activation in approximately 81% of AGRP/NPY neurons (25/31) with a rapid onset and a latency of 5.12 millisecond (**Fig. 5c-e**) indeed suggesting that BNC2 neurons directly inhibit AGRP/NPY neurons. Consistent with this, the oIPSCs were abolished after the application of the sodium channel blocker tetrodotoxin (TTX), which blocks action potential in all neurons, but inhibitory currents onto AGRP/NPY neurons were restored by the co-application of 4-aminopyride (4-AP), a potassium channel antagonist that enables ChR2-mediated depolarization of local presynaptic terminals (**Fig. 5c,f**). Finally, the oIPSCs observed in the presence of TTX and 4-AP were blocked after the application of the GABAA receptor antagonist picrotoxin (PTX) (**Fig. 5c,f**). In aggregate, these data show that BNC2 neurons directly inhibit AGRP/NPY neurons via the GABAA receptor.

**Fig. 5:**
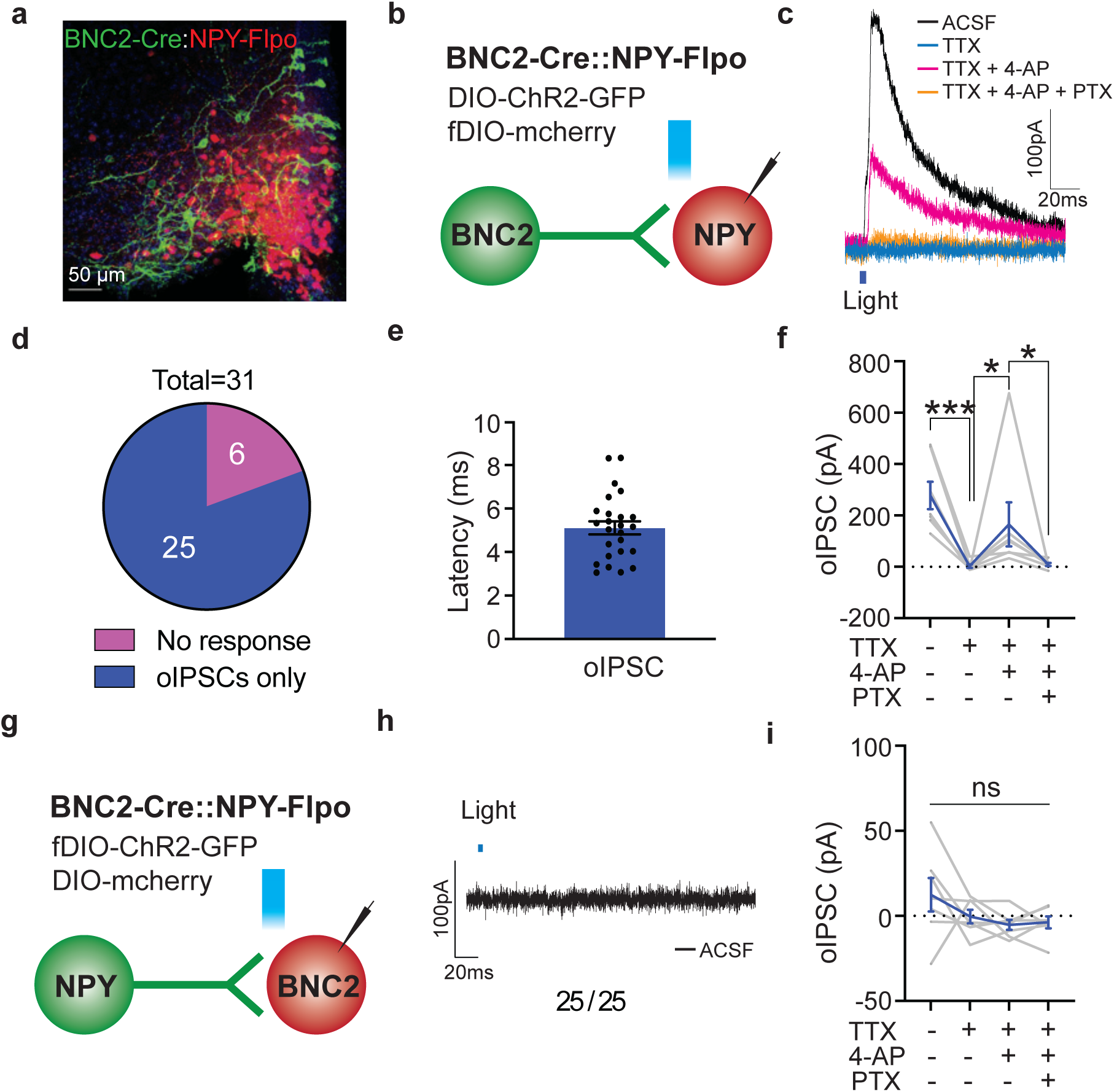
BNC2 neurons monosynaptically inhibit NPY neurons. **a**, Immunostaining of GFP and mCherry in ARC from adult BNC2-Cre::NPY-Flpo mice injected with DIO-ChR2-GFP and fDIO-mCherry. Scale bar, 50 µm. **b,g,** Schematics of ChR2-assisted circuity mapping of the connections between BNC2 neurons and NPY neurons. **c,** Representative oIPSCs from NPY neurons following activation of BNC2 neurons with different blockers. **d,** Percentage of responsive NPY neurons (oIPSCs detected) following BNC2 neuron activation. **e,** oIPSC latency upon light stimulation (N = 29 cells from 4 mice). **f,** Quantification of oIPSC amplitude in c (N = 7 cells from 4 mice). **h,** Representative oIPSCs from BNC2 neurons following activation of NPY neurons. **i,** Quantification of oIPSC amplitudes in different conditions (N = 7 cells from 4 mice). ns, not significant; *p<0.05; ***p<0.001. Details of the statistical analysis are provided as Source Data.

We also tested whether AGRP/NPY neurons could inhibit BNC2 neurons by injecting AAV-fDIO-ChR2-GFP and AAV-DIO-mCherry into the ARC of *BNC2-Cre*::*NPY-FlpO* mice. However, in contrast to the effects of BNC2 activation to increase oIPSCs in AGRP/NPY neurons, we failed to see an effect of AGRP/NPY activation on post-synaptic currents in mCherry labeled BNC2 neurons (0 out of 25) (**Fig. 5g-i**). This was despite the fact that brief blue light pulses triggered action potentials in GFP-labeled AGRP/NPY neurons (**Extended Data Fig. 6**). Thus while BNC2 neurons directly inhibit AGRP/NPY neurons, AGRP/NPY neurons do not alter the activity BNC2 neurons.

Previous reports have shown that AGRP/NPY neuronal activity is inhibited by the presence of food^21,35^. This raised the possibility that BNC2 neuronal activation might mediate some or all of the effect of food cues to inhibit AGRP/NPY neurons. We tested this by injecting an inhibitory DREADD, AAV-DIO-hM4Di-mCherry, together with an AAV-fDIO-GCamp6s into the ARC of *BNC2-Cre*::*NPY-FlpO* (**Fig. 6a**). After an overnight fast, we administered PBS or CNO 30 minutes prior to providing the animal with chow (**Fig. 6b**). Consistent with the prior results (**Fig. 4**), inhibition of BNC2 neurons significantly increased food intake relative to PBS-treated controls (**Fig. 6c**). We then imaged the activity of AGRP/NPY neurons after refeeding using fiber photometry with or without inhibition of BNC2 neurons. As previously reported, control mice receiving PBS showed a robust reduction in calcium signals in AGRP/NPY neurons when given food (**Fig. 6d,e**). We then measured the activity of AGRP/NPY neurons after giving CNO and found that the inhibition of BNC2 neurons significantly attenuated the decrease in AGRP/NPY activity seen after refeeding (**Fig. 6d,e**). These findings suggest that a portion of the sensory inputs that suppress AGRP/NPY neurons after refeeding are conveyed by BNC2 neurons.

**Fig. 6:**
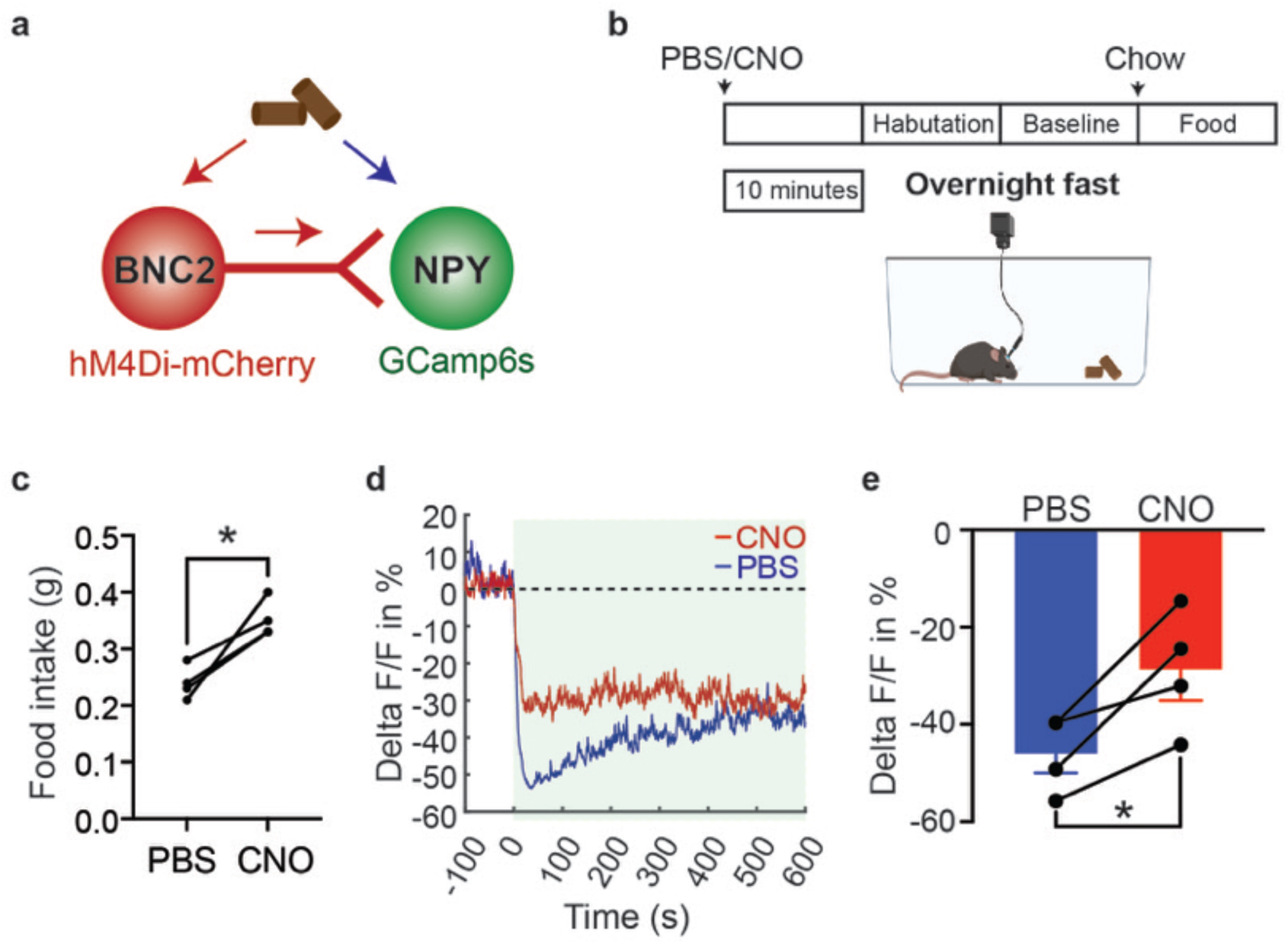
BNC2 neurons mediate the regulation of food cues on NPY neuron activity. **a**, The experimental design and hypothesis. **b,** Schematic of PBS or CNO administration time during fiber photometry recordings of NPY neuron activity. **c,** Food consumption during the 1O minutes after food presentation in mice injected with PBS or CNO (N = 4 mice). **d,** Average traces of calcium signals from NPY neurons aligned to the time of presentation of chow in mice receiving PBS (blue) or CNO (red). **e,** Quantification of fluorescence changes within 0-200 second timeframe in d (N = 4 mice). Data are presented as mean ± SEM. ns, not significant; *p<0.05. Details of the statistical analysis are provided as Source Data.

### Deletion of leptin receptors in BNC2 neurons causes hyperphagia and obesity

BNC2 neurons express the leptin receptor (**Fig. 1**) and are activated by leptin (**Fig. 2**) suggesting that they mediate some of leptin’s effects. We assess this by knocking out LepR in BNC2 neurons using CRISPR-Cas9 genome editing technology. BNC2-Cre mice were bred to LSL-Cas9-GFP mice^39^, and an AAV carrying two single guide RNAs (sgRNAs) designed to target the mouse *Lepr* (sgLepr) locus, or two control guide RNAs (sgCtrl)^23,40^, were injected bilaterally into the ARC. The specific deletion of the *Lepr* gene was confirmed by showing that leptin no longer increased the levels of pSTAT3 in BNC2 neurons that received injections of the sgLepr guide RNAs compared to those with sgCtrl guide RNAs (**Fig. 2, Extended Data Fig. 7**).

Mice with a LepR knockout in BNC2 neurons, gained significantly more weight compared to the sgCtrl-injected mice (28.14±0.37g and 35.57±1.43g for sgCtrl and sgLepr, respectively; p=0.0001, **Fig. 7a**). 8-weeks after the guide RNA injection, fat mass within the sgLepr group was also significantly increased relative to the sgCtrl group with a small increase in lean mass (**Fig. 7b**). Daily food intake was significantly higher in sgLepr mice at week 8, compared to their sgCtrl counterparts (4.27±0.21g and 5.24±0.28g for sgCtrl and sgLepr, respectively; p=0.0185, **Fig. 7c**). Total energy expenditure (EE) was significantly increased in the BNC2 LepR knockout mice and there was still a significant increase when it was indexed to lean mass (**Fig. 7d,e, Extended Data Fig. 8a**)^41^. However, this difference was no longer significant when the regression analysis of EE was calculated relative to body weight (**Extended Data Fig. 8b**). While the basis for the increased expenditure is not known, this indicates that the primary effect of the knockout to increase weight is a result of increased food intake. There was no difference in locomotor activity between the two groups (**Extended Data Fig. 8c,d**). The respiratory exchange ratio (RER) was higher in the sgLepr group in comparison to the sgCtrl group (**Extended Data Fig. 8e,f**), suggesting that there is increased carbohydrate utilization. Finally, we assayed glucose tolerance and found that sgLepr mice had increased fasting blood glucose levels (**Fig. 7f**). The BNC2 LepR knockout mice also showed impaired glucose tolerance during a glucose tolerance test (GTT), and a significantly reduced response to insulin compared to sgCtrl mice as demonstrated by the insulin tolerance test (ITT) (**Fig. 7g,h**). While the impaired glucose tolerance is consistent with the increased adiposity of the knockouts, the data do not exclude the possibility that these neurons directly regulate glucose metabolism. Taken together, these findings show that leptin signaling to ARC BNC2 plays a functional role to regulate food intake and body weight.

**Fig. 7:**
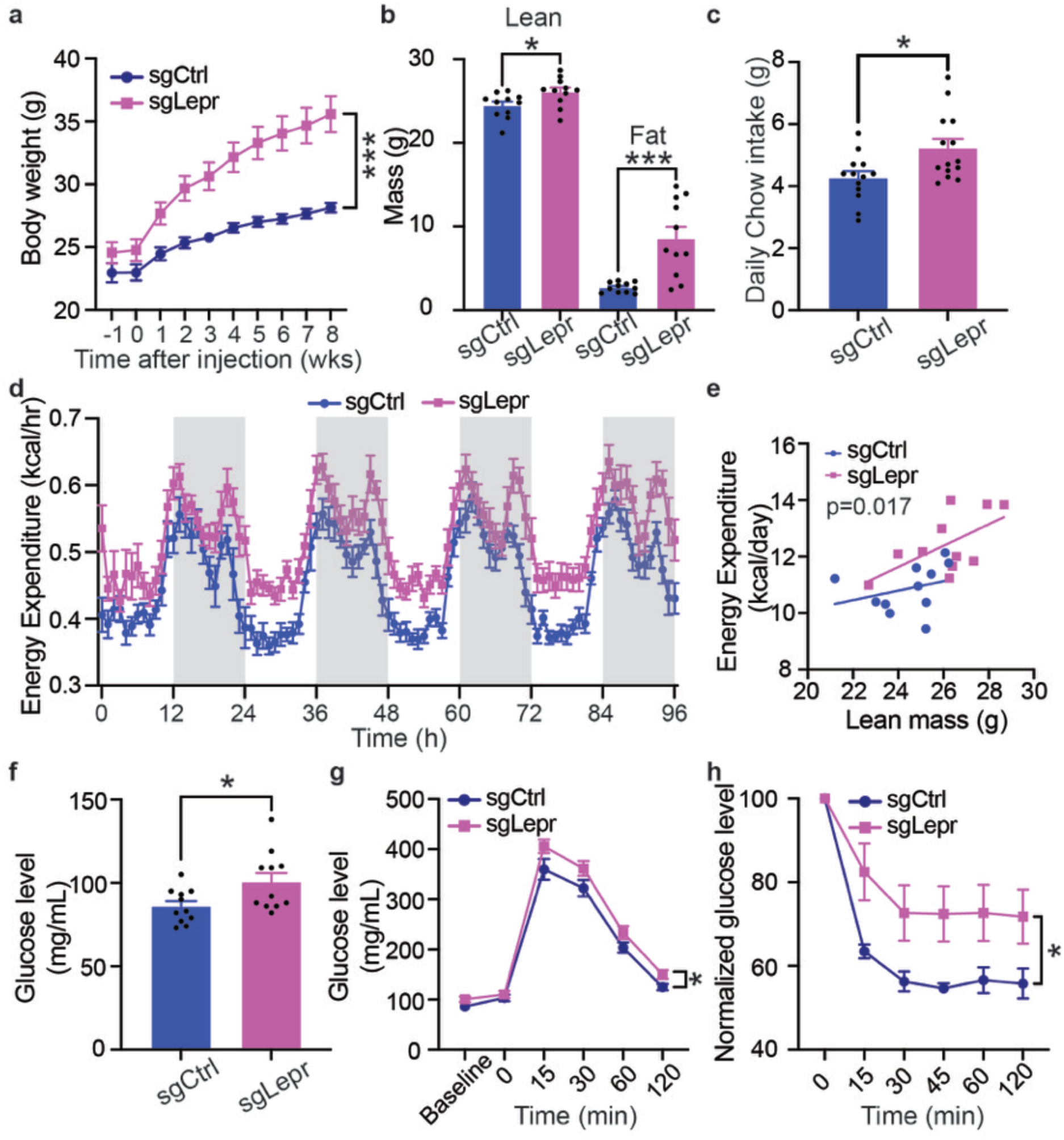
LepR knockout in BNC2 neurons causes hyperphagia and obesity. **a**, Body weight of mice fed on chow following injection of sgCtrl or sgLepr (N = 14 mice per group). **b,** Body composition of two groups of mice 8 weeks after virus injection (N = 11 mice per group). **c,** Daily chow intake of two groups at 8 weeks after virus injection (N = 13 sgCtrl-injected mice, N = 14 sgLepr-injected mice). **d,** Energy expenditure (EE) of two groups measured by indirect calorimetry at 10-12 weeks after virus injection. **e,** Regression analysis of EE vs. lean mass from d (N = 11 mice per group). **f,** Glucose levels of two groups after 16 h overnight fasting (N = 11 mice per group). **g,** Glucose tolerance test (GTT) of the two groups of mice at 10-12 weeks post virus injection (N = 11 mice per group). **h,** Insulin tolerance test (ITT) of the two groups of mice at 10-12 weeks post virus injection (N = 6 mice per group). Data are presented as mean ± SEM. ns, not significant; *p<0.05; ***p<0.001. Details of the statistical analysis are provided as Source Data.

## Discussion

Leptin is the afferent signal in a negative feedback loop regulating food intake and body weight. Leptin regulates the activity of discrete neural populations expressing LepR in the ARC and elsewhere^1^. In the ARC, leptin decreases the activity of orexigenic AGRP/NPY neurons and activates anorexigenic neurons expressing POMC. However, several lines of evidence have raised the possibility that there are additional, as yet unknown leptin-regulated neurons that suppress appetite (see below)^13,14,42^. However, the nature of this putative population(s) was not known.

Here, we report the use of snRNA-seq to identify a novel LepR-expressing neuron cluster in the ARC that expresses the *Bnc2* marker gene. While there are fewer BNC2/LepR neurons compared to those expressing POMC and AGRP, the level of LepR in BNC2 neurons is as high or higher than in these other populations. The data further show that BNC2 neurons are rapidly activated by leptin or the presence of food and that the level of activation is further modulated by food palatability and nutritional status (**Extended Data Fig. 9**). BNC2/LepR neurons directly inhibit AGRP/NPY neurons and BNC2 activation reduces food intake for a sustained period. Finally, in contrast to anorexigenic POMC neurons, a BNC2-specific knockout of LepR causes significant hyperphagia and obesity. These findings establish a role for BNC2 neurons to acutely induce satiety and thus add an important new cellular component to the neural pathways that maintain homeostatic control of energy balance.

The *Bnc2* gene encodes a conserved transcription factor that has been reported to play a role in cell development, proliferation, pigmentation, and liver fibrosis^43–46^. In support of a potential role to regulate weight, genome-wide association studies (GWAS) have consistently linked the *Bnc2* gene to body mass index, fat distribution, and diabetes^47–49^, although the underlying mechanisms have not been previously explored. This together with the data reported here suggests that *Bnc2* itself may play a role in the development or function of the BNC2/LepR neurons we identified. Further studies knocking out the *Bnc2* gene exclusively in BNC2/LepR neurons will be necessary to assess this possibility, currently underway. While the function of BNC2 in hypothalamus has not been ascertained, these studies establish that it is a highly specific marker for a novel population of cells expressing LepR. While LepR clusters not expressing AGRP/NPY or POMC have been previously identified in other studies, TRH, the canonical marker expressed in this cluster is also expressed in non-LepR clusters^26^. In contrast, in hypothalamus BNC2 expression is restricted to LepR neurons making it an especially useful marker for probing the function of the neurons in this cluster. This enabled us to establish an important role for these neurons to regulate food intake and mediate leptin action.

The canonical “yin-yang” model of feeding regulation has emphasized the opposing roles of POMC neurons (suppressing hunger) and AGRP/NPY neurons (promoting hunger)^2,10^. However, the characteristics of these neural populations are markedly different in several respects. For instance, while acute activation of AGRP/NPY neurons, using either opto- or chemo-genetic methods, rapidly induces feeding within minutes^3,4^, POMC neurons function as slow-acting satiety regulators, and opto- and chemo-genetic stimulation of POMC neurons only has a minimal effect on acute food intake^4,5,50^. Furthermore, an AGRP-specific LepR knockout in adult mice results in severe obesity but a knockout of LepR in adult POMC neurons has a significantly smaller effect on food intake and body weight^23^. In contrast, mutations that alter melanocortin signaling including POMC, the PCSK1/2 processing enzymes, and MC4R cause severe obesity while mutations that alter signaling in AGRP/NPY neurons have not been reported to alter weight^8,15,51^. Finally, while leptin injections acutely suppress food intake, particularly in ob/ob mice, the activity of AGRP/NPY and POMC neurons is not significantly altered by acute leptin injection^11,12^. In aggregate, these findings raise the possibility that there might be a missing leptin-regulated neuronal population that acutely suppresses food intake. Consistent with this, a knockout of LepR in GABAergic neurons had a greater effect than did a knockout of LepR in AGRP or POMC neurons^13^. The identity of this “missing” GABAergic population was not known and the evidence reported here showing that BNC2/LepR neurons rapidly suppress feeding suggests that they serve this function. However, these data do not exclude the possibility that additional populations might also contribute.

Optogenetic activation of BNC2 neurons potently reduces food intake for a sustained period even after photoactivation has ceased. This is similar to the sustained effect of AGRP/NPY neuronal activation which induces a sustained increase in food intake through the release of neuropeptide Y^52,53^. The sustained satiety effect lasted at least 20 minutes and could potentially persist even longer. Previous research indicates that sustained hunger after AGRP/NPY neuron activation can extend up to 1 hour^52^. Furthermore, the duration of the satiety effect may be influenced by the duration of BNC2 neuron activation, akin to the pattern observed with AGRP/NPY neuron activation^52^. BNC2 neurons are GABAergic and directly inhibit AGRP/NPY neurons via the GABAA receptor though, as is the case for other feeding neurons, it is possible that neuropeptides also contribute. The snRNA-seq data shows expression of several putative neuropeptides in BNC2 neurons including *Scg2, Chga,* and *Trh,* which have been implicated in feeding control and energy balance^54–59^. Future studies will be necessary to determine whether in addition to GABA, these neuropeptides mediate some of the effects of BNC2 neurons.

BNC2 neurons suppress appetite, at least in part, by directly inhibiting AGRP/NPY neurons via monosynaptic connections. However, similar to POMC neurons, this interaction is not reciprocal^60,61^ as AGRP/NPY neurons do not appear to modulate the activity of BNC2 neurons. Similar to AGRP/NPY neurons, BNC2 neurons are activated by sensory cues associated with food. Thus, similar to AGRP/NPY neurons BNC2 neurons function as coincidence detectors for both sensory and interoceptive inputs such as leptin. The rapid regulation of AGRP/NPY and BNC2 neurons by food cues is likely mediated by associative sensory inputs, the precise source of which is currently unknown. It is plausible that these sensory inputs synapse onto both BNC2 neurons and AGRP/NPY neurons, exerting distinct effects on these two neuronal populations. If true, mapping the afferent inputs to both may reveal anatomic sites that convey associative sensory inputs. Alternatively, the food cues may primarily target BNC2 neurons, which subsequently relay the information to AGRP/NPY neurons. In support of the former, we find that inhibiting BNC2 neurons only partially blunts the effect of food cues to suppress AGRP/NPY activity suggesting both direct and indirect (via BNC2 neurons) pathways by which sensory inputs suppress AGRP/NPY neurons. It has also been shown that while food cues can transiently suppress AGRP/NPY activity, caloric content and consumption are required for a sustained reduction in their activity^62^. This is also the case for BNC2 neurons because the level of BNC2 neuron activity is markedly reduced when the food is not accessible.

Conditions of energy deficit activate AGRP/NPY neurons eliciting the aversive feeling of hunger. A recent study revealed that AGRP/NPY neurons stimulate appetite by “teaching” animals that food or food cues can suppress the negative valence associated with hunger^63^. The rapid reduction in the aversive effect of activating AGRP/NPY neurons by food cues eventually enhances the incentive salience of these cues, thereby facilitating learning of food acquisition tasks over time. Similar to AGRP/NPY neurons, the level of activation of BNC2 neurons is dependent on food availability, the duration of its presence, food palatability, and energy state but in contrast, BNC2 neuron activation is associated with positive valence. However, this response is only evident after fasting when AGRP/NPY activity is high. This suggests that BNC2 activation contributes to these learned responses and that food can both induce reward and suppress negative valence in fasted mice.

Deletion of LepR in adult BNC2 neurons resulted in abnormal GTTs and ITTs, implying a potential regulatory role of BNC2 neurons in glucose metabolism. The impaired glucose tolerance and insulin action in BNC2 LepR knockout mice could be a secondary consequence of the pronounced obesity that develops. However, it is also possible that BNC2 neurons may directly fluence on glucose metabolism similar to the acute activation of AGRP/NPY neurons which impairs systemic insulin sensitivity, independent of its effect to increase feeding^24^. Here again, some of these effects could be a result of the effect of BNC2 neurons on AGRP/NPY neural activity. We also found that despite increasing food intake and body weight, a LepR knockout in BNC2 neurons paradoxically increased energy expenditure though this was no longer significant when it was corrected for body weight. This is in contrast to a global knockout of LepR in db/db mice which show decreased energy expenditure and also reduced lean^64,65^. The basis for the increased lean mass and EE in the BNC2 LepR knockouts is not known and suggests that other neural pathways contribute to the reduced energy expenditure and lower lean mass associated with defects in leptin signaling. The identification of the specific pathways responsible for this will require further investigation as will an explanation for the increased energy expenditure in the BNC2 LepR knockouts. In aggregate, the data suggest that the increased weight of the BNC2 LepR knockouts is a result of hyperphagia.

In summary, we have identified BNC2 neurons as a fast-acting population of neurons that acutely regulate feeding and energy balance bidirectionally. These findings add an important new component to the neural circuit that regulates appetite and adiposity, while also shedding new light on the mechanisms by which leptin regulates this system. Finally, pharmacologic activation of these neurons could have therapeutic implications to reduce weight or to suppress the negative valence associated with hunger^66^.

## Methods

### Mice

All animal experiments were approved by the Institutional Animal Care and Use Committee at Rockefeller University and were carried out in accordance with the National Institutes of Health guidelines. Mice were group housed in a 12-h light/12-h dark cycle at 22 °C and 30–60% humidity with ad-libitum access to a regular chow diet and water. We used the following genotypes of mice: C57BL/6J (wild type; number 000664, The Jackson Laboratory), NPY-IRES2-FlpO-D (The Jackson Laboratory, stock no. 030211), Rosa26-LSL-Cas9 (The Jackson Laboratory, stock no. 024857), BNC2-P2A-iCre (made in-house). For all Cre or Flp mouse line experiments, only heterozygous animals were used. Sample sizes were decided based on experiments from similar studies. Littermates of the same sex were randomly assigned to either experimental or control groups.

### Generation of BNC2-P2A-iCre mouse line

The BNC2-P2A-iCre mouse line was generated by the CRISPR and Genome Editing Resource Center and Transgenic and Reproductive Technology Resource Center at Rockefeller University using CRISPR-Cas9 technology^29^. Briefly, a custom-designed long single-stranded DNA (lssDNA) donor, containing homology arms of *Bnc2* locus flanking the P2A-iCre sequence, was inserted near the endogenous STOP codon. Two guide RNAs (sgRNAs) were employed to induce site-specific double-stranded breaks. The lssDNA donor with the pre-assembled Cas9 protein/gRNA complexes was mixed and microinjected into C57BL/6J (The Jackson Laboratory) mouse zygotes following standard CRISPR genome engineering protocols. The resulting live offspring were genotyped through PCR with two sets of primers that specifically amplified the mutant allele. Validation was ensured via Sanger sequencing. The BNC2-P2A-iCre transgenic animals were bred to C57BL/6J mice for maintenance.

### Viruses

AAV viruses used in these studies were obtained from Addgene, UNC Vector Core, or generated through Janelia Viral Tools Service. We used the following viruses: AAV5-hSyn-DIO-hM3D(Gq)-mCherry (Addgene, #44361, 2.2 × 10^13^ vg/mL), AAV5-hSyn-DIO-hM4Di(Gi)-mCherry (Addgene, #44362, 2.5 × 10^13^ vg/mL), AAV5-hSyn-DIO-mCherry (Addgene, #50459, 2.2 × 10^13^ vg/mL), AAV5-Ef1a-DIO-EYFP (Addgene, #27056, 1.6 × 10^13^ vg/mL), AAV5-hSyn-Flex-GCaMP6s-WPRE (Addgene, #100845, 2.9 × 10^13^ vg/mL), AAV5-EF1a-DIO-hChR2(H134R)-EYFP (UNC Vector Core, 2.7 × 10^12^ vg/mL), AAV1-hSyn1-SIO-stGtACR2-FusionRed (Addgene, #105677, 2.1 × 10^13^ vg/mL), AAV5-Ef1a-fDIO-mCherry (Addgene, #114471, 2.3 × 10^13^ vg/mL), AAV8-Ef1a-fDIO-GCamp6s (Addgene, #105714, 2.1 × 10^13^ vg/mL), AAV5&DJ-EF1a-fDIO-hChR2(H134R)-EYFP-WPRE (UNC Vector Core, 1.4 × 10^12^ vg/mL). For *Lepr* deletion, AAV viral vectors were cloned in-house and packaged with the AAV5 serotype through Janelia Viral Tools Service. The sequences of sgLepR are: 5’-GAGTCATCGGTTGTGTTCGG-3’, 5’-TGCCGGCGGTTGGATG GACT-3’ (virus titer, 4.9 × 10^12^ vg/mL); The sequence of sgCtrl is: 5’-TTTTTTTTTTTTTTGAATTC-3’ (virus titer, 8.5 × 10^12^ vg/mL). Viral aliquots were stored at −80 °C before stereotaxic injection.

### Stereotactic surgery

Mice (8–10 weeks old) were anesthetized using isoflurane anesthesia (induction 5%, maintenance 1.5–2%) and positioned on a stereotaxic rig (Kopf Instruments, Model 1900). Viruses were delivered into the brains through a glass capillary using a Drummond Scientific Nanoject III Programmable Nanoliter Injector. For the ARC region, the following coordinates relative to the bregma were used: anterior-posterior (AP), −1.65 mm to −1.70 mm; medial-lateral (ML), ± 0.25 mm to 0.30 mm; and dorsal-ventral, (DV) −5.9 mm. For chemogenetics experiments, Bnc2 neuron labeling, and *Lepr* deletion, 30-50 nl of the virus was injected bilaterally at a rate of 1 nl/s. For optogenetics, 30 nl of the virus was injected unilaterally at a rate of 1 nl/s followed by the implant of an optical fiber (ThorLabs, CFM12U-20) at 200 µm above the ARC (AP: −1.65 mm, ML: 0.3 mm, DV: −5.7 mm). For fiber photometry experiments, 30 nl of the virus was injected unilaterally followed by the implant of an optical fiber cannula (Doric, MFP_400/430/1100-0.57_1m_FCM-MF2.5_LAF) at 150 µm above the ARC (AP: −1.65 mm, ML: 0.3 mm, DV: −5.75 mm). For CRACM experiments, the two viruses were mixed at the ratio of 1:1, and 50 nl of the mixed virus was injected bilaterally into the ARC.

### Isolation of nuclei and snRNA-seq

Male C57BL/6J mice aged 10-12 weeks were euthanized via transcardial perfusion using ice-cold HEPES-Sucrose Cutting Solution containing NaCl (110 mM), HEPES (10 mM), glucose (25 mM), sucrose (75 mM), MgCl2 (7.5 mM), and KCl (2.5 mM) at a PH of 7.4. Subsequently, brains were quickly dissected in the same solution, frozen using liquid nitrogen, and stored at −80 °C until nuclei isolation. To isolate nuclei, as previously described, the samples were thawed on ice, resuspended in HD buffer containing tricine KOH (10 mM), KCl (25 mM), MgCl2 (5 mM), sucrose (250 mM), 0.1% Triton X-100, SuperRNaseIn (0.5 U/ml), RNase Inhibitor (0.5 U/ml). Homogenization was performed using a 1-ml dounce homogenizer. The resulting homogenates were filtered using a 40 μM filter, centrifuged at 600g for 10 minutes, and resuspended in nucleus storage buffer containing sucrose (166.5 mM), MgCl2 (10 mM), Tris buffer (pH 8.0, 10 mM), SuperRNaseIn (0.05 U/ml), RNase Inhibitor (0.05 U/ml) for subsequent staining. Nucleus quality and number were assessed using an automated cell counter (Countess II, Thermo Fisher). For staining, nuclei were labeled with Hoechst 33342 (Thermo Scientific H3570; 0.5 µl per million nuclei), anti-NeuN Alexa Fluor 647-conjugated antibody (Abcam ab190565) (0.5 µl per million nuclei), and TotalSeq anti-Nuclear Pore Complex Proteins Hashtag antibody (BioLegend 682205) (0.5 mg per million nuclei) for 15 minutes at 4 °C. Following staining, samples were washed with 10 ml 2% BSA (in PBS) and centrifuged at 600g for 5 minutes. Nuclei were then resuspended in 2% BSA (in PBS) with RNase inhibitors (SuperRNaseIn 0.5 U/ml, RNase Inhibitor 0.5 U/ml) for subsequent fluorescence-activated cell sorting. The samples were gated based on Hoechst fluorescence to identify nuclei and then further sorted based on high Alexa Fluor 647 expression, designating NeuN+ nuclei as neurons.

Nuclei were captured and barcoded using 10x Genomics Chromium v3 following the manufacturer’s protocol. The processing and library preparation were carried out by the Genomics Resource Center at Rockefeller University, and sequencing was performed by Genewiz using Illumina sequencers.

### SnRNA-seq analysis

The FASTQ file was analyzed with Cell Ranger version 5.0. The snRNA-seq data for ARC (WT) was preprocessed individually using the Seurat v4 (v4.0.3). Cells with more than 800 and fewer than 5,000 RNA features were selected for further analysis. Cells with greater than 1% mitochondrial genes and greater than 12,000 total RNA counts were also removed. Genes detected in fewer than 3 cells were excluded. Then we demultiplexed the cells based on their hashtag count (positive. quantile =0.8) using the built-in function in Seurat v4. Only the cells with singlet Hashtag assignment were kept for downstream analysis. The data was then log-normalized with a scale factor of 10,000. After the initial quality control, demultiplexing, and normalization steps, all the singlets were then scaled, and dimensionally reduced with PCA and UMAP. Leiden clustering (resolution =0.55) was applied to identify clusters. We used known cell-type specific gene expression to annotate the clusters. The analytic code and processed data are available upon request.

Co-expression analysis of marker genes within the human ARC was performed using previously published human adult samples and the data can be accessed through the NeMO archive (https://assets.nemoarchive.org/dat-917e9vs). A cell was considered to express the marker gene if at least 2 unique molecular identifiers (UMIs) were detected. The identification of arcuate cells was achieved through clustering and the expression of canonical markers, as detailed in the earlier study. Co-expression of genes like *Lepr*, *Bnc2*, *Agrp*, *Npy*, and *Pomc* was tabulated in R, and two-tailed Fisher’s tests were calculated to assess the significance of co-expression of gene pairs within the 16,819 arcuate cells in the human dataset.

### Chemogenetics for activation or inhibition

AAV viruses were bilaterally delivered into the ARC of male BNC2-iCre mice aged 8-10 weeks. Mice were then allowed to recover and express DREADDs for at least 3 weeks. For activation or inhibition, animals were intraperitoneally injected with 3 mg/kg of CNO or PBS (control).

### Optogenetics for activation or inhibition

AAV viruses were unilaterally delivered into the ARC of male BNC2-iCre mice aged 8-10 weeks followed by the implantation of an optic fiber. Subsequently, the mice were given a recovery period of at least 3 weeks to allow for gene expression. Prior to the experiments, the mice were habituated to patch cables over a period of 5 days. The implanted optic fibers were connected to patch cables using ceramic sleeves (Thorlabs) and linked to a 473 nm laser (OEM Lasers/OptoEngine, Midvale, UT). The output of the laser was verified at the beginning of each experiment. A blue light, generated by a 473 nm laser diode (OEM Lasers/OptoEngine) with a power of 15 mW, was used. The light pulse (10 ms) and frequency (20 Hz) were controlled through a waveform generator (Keysight) to either activate or inhibit Bnc2 neurons in the ARC. In the activation feeding experiments, mice were allowed to acclimate to the cage for 20 minutes. Subsequently, three feeding sessions, each lasting 20 minutes, were initiated. During these sessions, the light was turned off for the initial 20 minutes, switched on for the subsequent 20 minutes, and then turned off again for the remaining 20 minutes. In the inhibition feeding experiments, following the 20-minute acclimation, each feeding session was extended to 30 minutes. The amount of food consumed during each feeding session was manually recorded. Animals were sacrificed at the end of the experiments to confirm viral expression and fiber placement using immunohistochemistry.

### Real-time place preference

A custom-made 2-chamber box (50 × 50 × 25 cm black plexiglass) with an 8.5 cm gap enabling animals to move freely between the chambers is used for this assay. To evaluate the mice’s initial preference, they were introduced into the box for a 10-minute session without any photostimulation. Subsequently, in the second 10-minute session following the initial one, photostimulation (15 mW, 20 Hz) was paired with the chamber that the mice exhibited less preference for during the initial session. The Ethovision software, coupled with a CCD camera, facilitated the recording of the percentage of time spent by the mice in each chamber.

### Fiber photometry

Mice were acclimated to tethering and a home cage-style arena for 5 minutes daily over the course of 5 days prior to the experiment. Data acquisition was conducted using a Fiber Photometry system from Tucker-Davis Technologies (RZ5P, Synapse) and Doric components, with recordings synchronized to video data in Ethovision through TTL triggering. A dual fluorescence Mini Cube (Doric) combined light from 465nm and isosbestic 405nm LEDs, which were transmitted via the recording fiber connected to the implant. GCaMP6s fluorescence, representing the calcium-dependent signal (525nm), and isosbestic control (430nm) were detected using femtowatt photoreceivers (Newport, 2151) and a lock-in amplifier at a sampling rate of 1kHz. Analysis was conducted using a Matlab script involving the removal of bleaching and movement artifacts through a polynomial least square fit applied to the 405nm signal, adjusting it to the 465nm trace (405fitted), and then calculating the GCaMP signal as % Δ F/F = (465signal-405fitted)/405fitted. The resulting traces were filtered using a moving average filter and down-sampled by a factor of 20. The code is available upon request.

### In situ hybridization

Mice were briefly transcardially perfused with ice-code RNase-free PBS. Brains were then quickly collected, embedded in OCT on dry ice, and stored at −80°C until cryostat sectioning (15 µm thickness) onto Superfrost Plus Adhesion Slides (Thermo Fisher). RNAscope Fluorescent Multiplex assay (Advanced Cell Diagnostics Bio) was performed based on the manufacturer’s protocol. All reagents were purchased from Advanced Cell Diagnostics (ACDbio). Probes for the following mRNAs were used: Agrp (cat no. 400711-C3), Pomc (cat no. 314081-C3), Lepr (cat no. 402731), Slc31a1 (cat no. 319191), Bnc2 (cat no. 518521-C2). Briefly, brain sections were fixed in 4% PFA at 4°C for 15 minutes followed by serial submersion in 50% EtOH, 70% EtOH, and twice in 100% EtOH for 5 minutes each at room temperature. Sections were treated with Protease IV for 30 minutes at RT followed by a 2-hour incubation with specific probes at 40°C using a HyBez oven. Signal amplification was achieved through successive incubations with Amp-1, Amp-2, Amp-3, and Amp-4 for 30, 15, 30, and 15 minutes, respectively, at 40°C using a HyBez oven. Each incubation step was followed by two 2-minute washes in RNAscope washing buffer. Nucleic acids were counterstained with DAPI Fluoromount-G (SouthernBiotech) mounting medium before coverslipping. The slides were visualized using an inverted Zeiss LSM 780 laser scanning confocal microscope with a ×20 lens. The acquired images were imported into Fiji for further analysis.

### Immunohistochemistry

Mice were transcardially perfused with PBS first and then 4% PFA for fixation. Brains were collected and immersed in 4% PFA overnight at 4°C for additional fixation. Fixed brains were sequentially immersed in 10% sucrose, 20% sucrose, and 30% sucrose buffers for 1 hour, 1 hour, and overnight, respectively, all at 4°C. After this, the brains were embedded in OCT and stored at −80°C until cryostat sectioning (30-50 µm thickness). For the staining process, brain sections were first blocked in a blocking buffer containing 3% BSA, 2% goat serum, and 0.1% triton X-100 in PBS for 30 minutes at room temperature followed by an overnight incubation with primary antibodies in the cold room. After washing in PBS, the sections were incubated with fluorescence-conjugated secondary antibodies (Invitrogen) for 1h at room temperature. Stained sections were mounted onto SuperFrost (Fisher Scientific 22-034-980) slides and then visualized with an inverted Zeiss LSM 780 laser scanning confocal microscope with a ×10 or ×20 lens. The acquired images were imported to Fiji for further analysis. The following antibodies were used: cFos antibody (1:1000, Synaptic systems, #226308), pSTAT3 antibody (1: 1000, Cell Signaling Technology, #9145s), GFP (1:1000, abcam, ab13970), RFP (1:1000, Rockland, #600-401-379).

### Electrophysiology and channelrhodopsin-assisted circuit mapping (CRACM)

Adult mice were euthanized via transcardial perfusion using ice-cold cutting solution containing choline chloride (110 mM), NaHCO3 (25 mM), KCl (2.5 mM), MgCl2 (7 mM), CaCl2 (0.5 mM), NaH2PO4 (1.25 mM), glucose (25 mM), ascorbic acid (11.6 mM), and pyruvic acid (3.1 mM). Subsequently, brains were quickly dissected in the same solution and sectioned through a vibratome into 275 µm coronal sections. These sections were then incubated in artificial cerebral spinal fluid containing NaCl (125 mM), KCl (2.5 mM), NaH2PO4 (1.25 mM), NaHCO3 (25 mM), MgCl2 (1 mM), CaCl2 (2 mM), and glucose (11 mM) at 34 °C for 30 minutes, followed by room temperature incubation until use. The intracellular solution for current-clamp recordings contained K-gluconate (145 mM), MgCl2 (2 mM), Na2ATP (2 mM), HEPES (10 mM), EGTA (0.2 mM, 286 mOsm, pH 7.2). The intracellular solution for the voltage clamp recording contained CsMeSO3 (135 mM), HEPES (10 mM), EGTA (1 mM), QX-314 (chloride salt, 3.3 mM), Mg-ATP (4 mM), Na-GTP (0.3 mM), and sodium phosphocreatine (8 mM, pH 7.3 adjusted with CsOH). Signals were acquired using the MultiClamp 700B amplifier and digitized at 20 kHz using DigiData1550B (Molecular Devices). The recorded electrophysiological data were analyzed using Clampfit (Molecular Devices) and MATLAB (Mathworks).

For CRACM experiments, voltage-clamp recordings were conducted on Bnc2 and Npy neurons. To record oIPSCs, the cell membrane potential was held at 0 mV. ChR2-expressing axons were activated using brief pulses of full-field illumination (0.5 ms, 0.1 Hz, 10 times) onto the recorded cell with a blue LED light (pE-300 white; CoolLED). Subsequently, TTX (1 µM), 4-AP (100 mM), and PTX (1 µM) were sequentially applied through the bath solution, each for 10–20 minutes. Data acquisition commenced at least 5 minutes after each drug application.

### Indirect calorimetry

Indirect calorimetry was performed using the Phenomaster automated home cage phenotyping system (TSE Systems, Bad Homburg, Germany). Mice were individually housed in environmentally controlled chambers maintained at 22°C, following a 12-h light/12-h dark, and at 40% humidity, with ad libitum access to food and water. O2 and CO2 measurements were collected at 15-minute intervals with a settling time of 3 minutes and a sample flow rate of 0.25 L/minute. The raw data obtained were analyzed using CalR^41^.

### Blood glucose, glucose, and insulin tolerance test

Blood glucose levels were measured using a OneTouch Ultra meter and glucose test strips. For glucose tolerance tests (GTT), mice were fasted overnight followed by a 20% glucose injection (2 g/kg) and glucose measurements at 0, 15, 30, 60, and 120 minutes. Insulin tolerance tests (ITT) were conducted after a 4-hour fast, with insulin injection (0.75 U/kg) and glucose measurements at 0, 15, 30, 45, 60, and 120 minutes.

### Statistical analysis

All statistical analyses were performed in GraphPad Prism 9. Data distribution was tested for normality (Shapiro-Wilk test) and then comparisons were made using parametric or non-parametric tests, as appropriate. Two-tailed statistical tests were used, and statistical significance was determined by Student’s t-test, Mann-Whitney test, Fisher’s exact test, one-way or 2-way ANOVA, and Friedman test as indicated in the Source Data.

## Data availability

All data generated or analyzed during this study are included in the manuscript and supporting files. The raw sequencing data have been deposited in the Gene Expression Omnibus and are available under the accession number GSE249564. The processed data are available upon request.

## Code availability

The analytic code is available upon request.

## Acknowledgments

We thank all members of the laboratory of J.M.F. for discussions and support, Hong Duan from the Genomics Resource Center, and Flow Cytometry Resource Center for technical assistance with snRNA-seq experiments; Chingwen Yang from CRISPR and Genome Editing Center for generating the BNC2-P2A-iCre mouse line; and Bio-Imaging Resource Center. This work was supported by the JPB Foundation and the Howard Hughes Medical Institute (to J.M.F.) and the Kavli Neural Systems Institute Postdoctoral Fellowship (to H.L.T).

## Author contributions

H.L.T. conceived, designed, performed, and analyzed the research with inputs from all authors.

L.Y. performed electrophysiology experiments and analyzed the data. Y.T. and P.W. analyzed the mouse snRNA-seq data. J.I., K.P., and A.I. performed experiments. B.R.H. analyzed the human snRNA-seq data. P.C. and D.L. provided inputs on experimental design. C.K. provided the code for fiber photometry data analysis. H.L.T. and J.M.F. wrote the manuscript.

## Materials & Correspondence

Correspondence to Jeffrey M. Friedman

---

**Extended Data Fig. 1:**
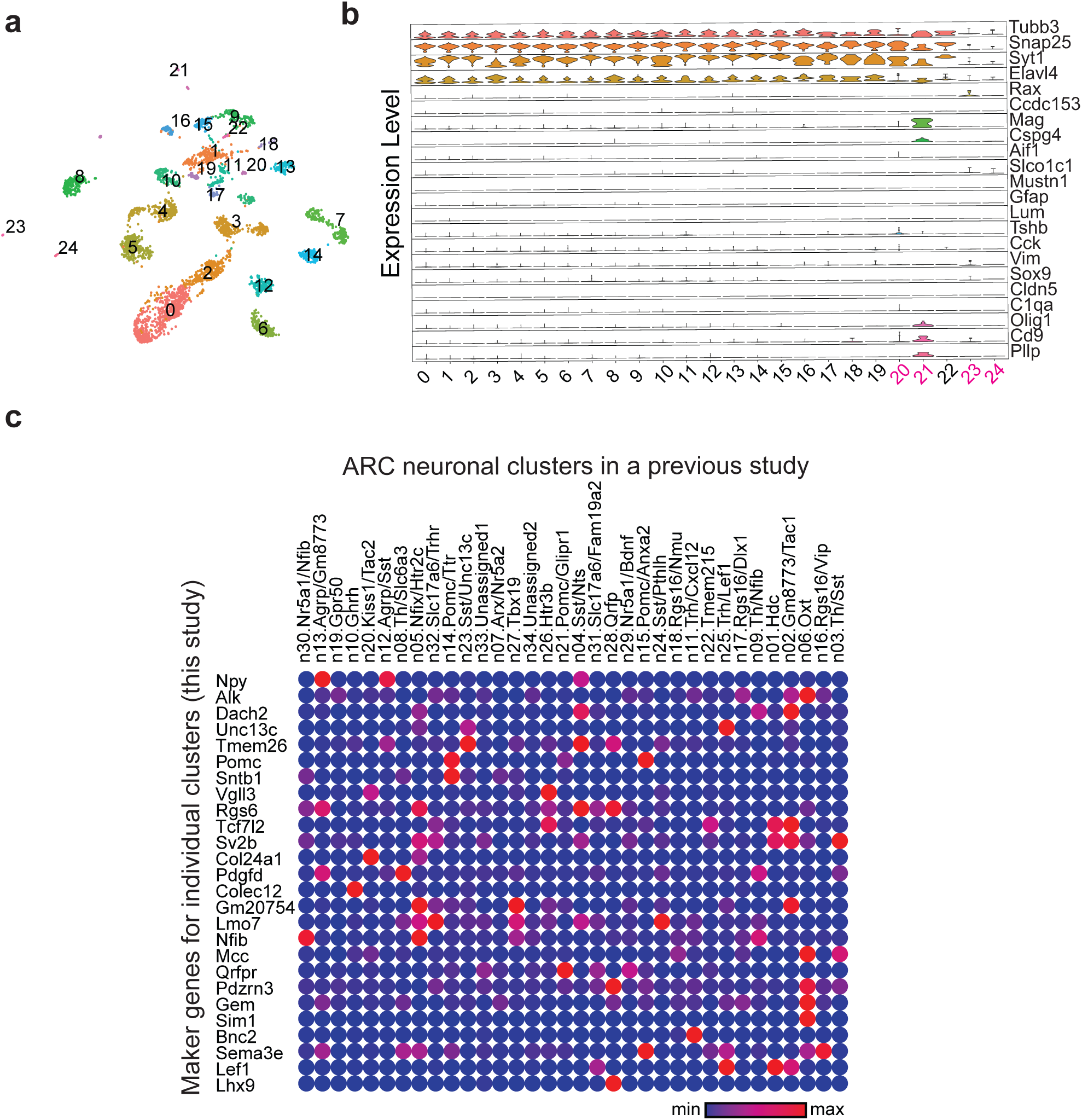
SnRNA-seq analysis of mouse ARC neurons. **a**, Single-cell UMAP plot of the ARC (N = 6 adult male WT mice). **b,** Violin plot of known markers of different cell types: neurons (Tubb3, Snap25, Syt1, Elavl4), tanycytes (Rax), ependymocytes (Ccdc153), oligodendrocyte cells (Mag, Olig1, Cspg4, Cd9, Pllp), macrophages (Aif1, C1qa), endothelial cells (Slco1c1, Cldn5), mural cells (Mustn1), astrocyte (Gfap, Sox9), vascular and leptomeningeal cells (Lum), pituitary cells (Tshb), pars tuberalis cells (Cck), ependymal cells (Vim). Clusters 20/21/23/24 are non-neuronal clusters. **c,** Dot plot displaying the expression of cluster-specific marker genes (identified in this study) across neuronal clusters with the ARC from a previous study26.

**Extended Data Fig. 2:**
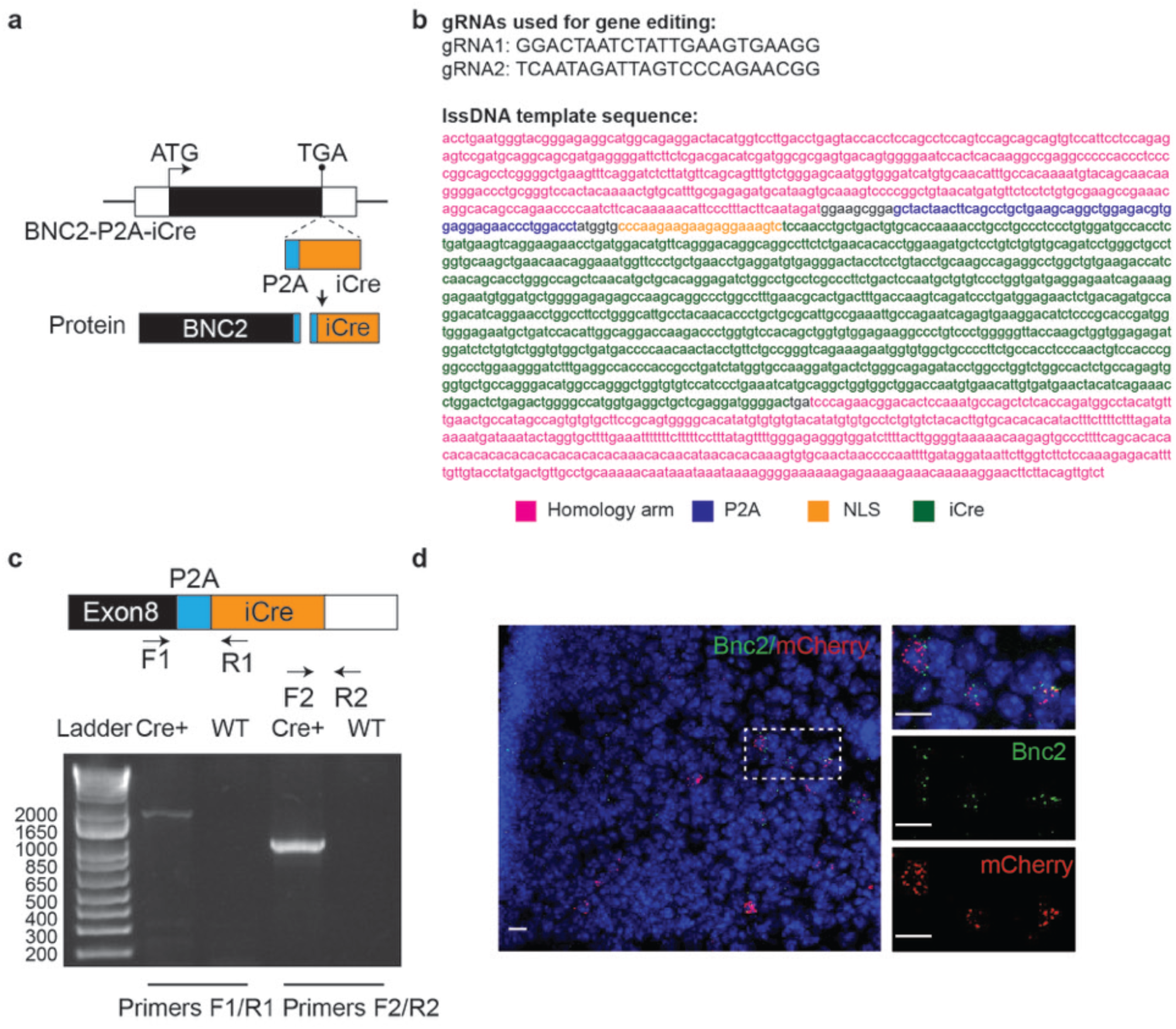
Generation of BNC2-P2A-iCre knockin mouse line. **a**, Schematic of BNC2-P2A-iCre locus. **b,** Sequence information of gRNAs and lssDNA. **c,** Genomic PCR using specific primers. **d,** Cre expression was marked using a Cre-dependent mCherry viral construct injection, and its colocalization with endogenous Bnc2 mRNA was confirmed through RNA ISH. Scale bar, 20 µm.

**Extended Data Fig. 3:**
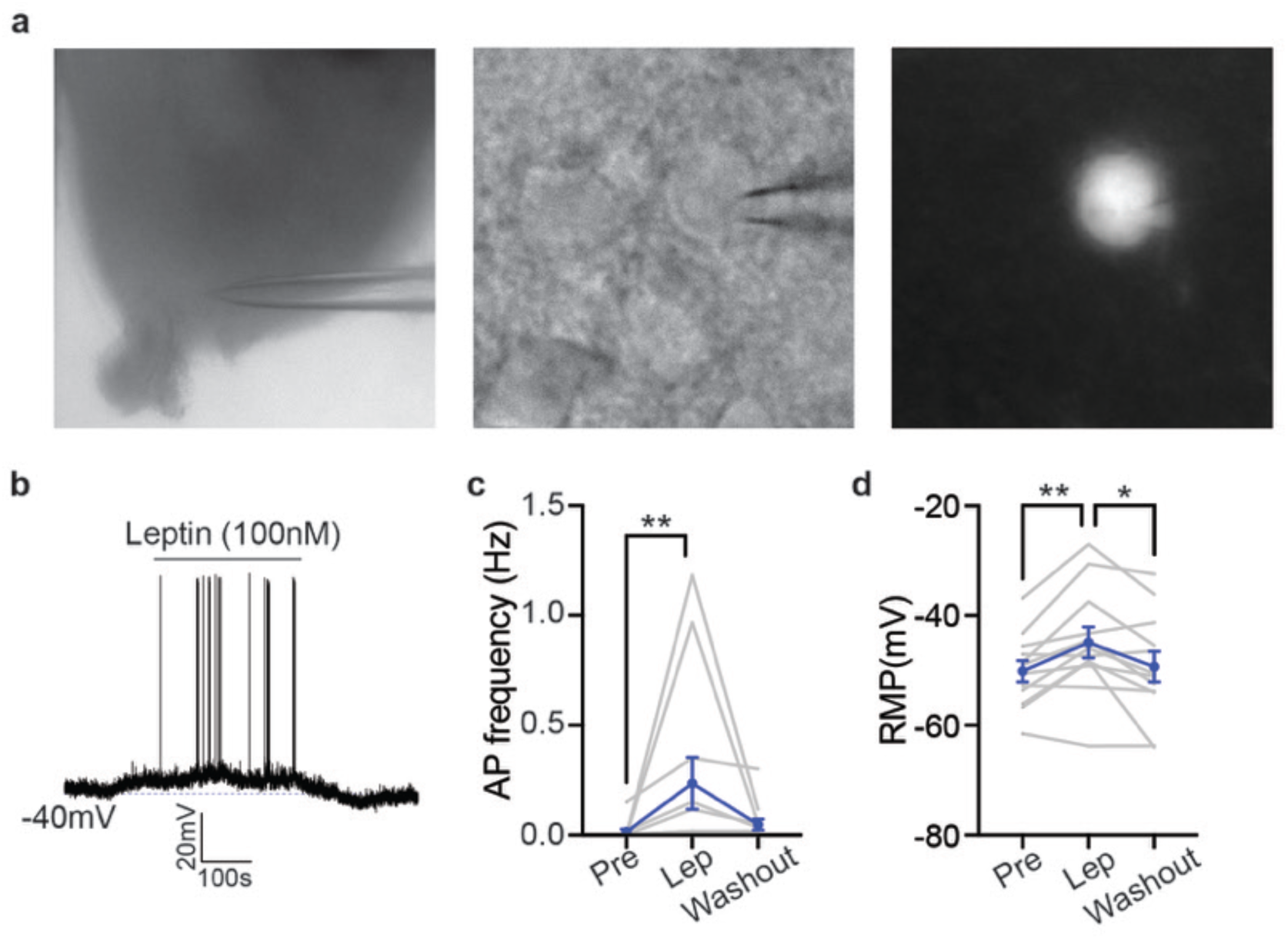
Activation of BNC2 neurons by leptin. **a**, Representative images of a patched GFP-labeled BNC2 neuron. **b,** Spontaneous APs of BNC2 neurons in ARC from ad-libitum-fed adult male BNC2-Cre mice (injected with DIO-GFP). Leptin (100 nM) was added to the bath solution at the indicated time window. **c,d,** Spontaneous AP frequency **(c)** and RMP **(d)** of BNC2 neurons before, during, and after leptin application (N = 12 cells from 4 mice). Data are presented as mean ± SEM. *p<0.05; **p<0.01. Details of the statistical analysis are provided as Source Data.

**Extended Data Fig. 4:**
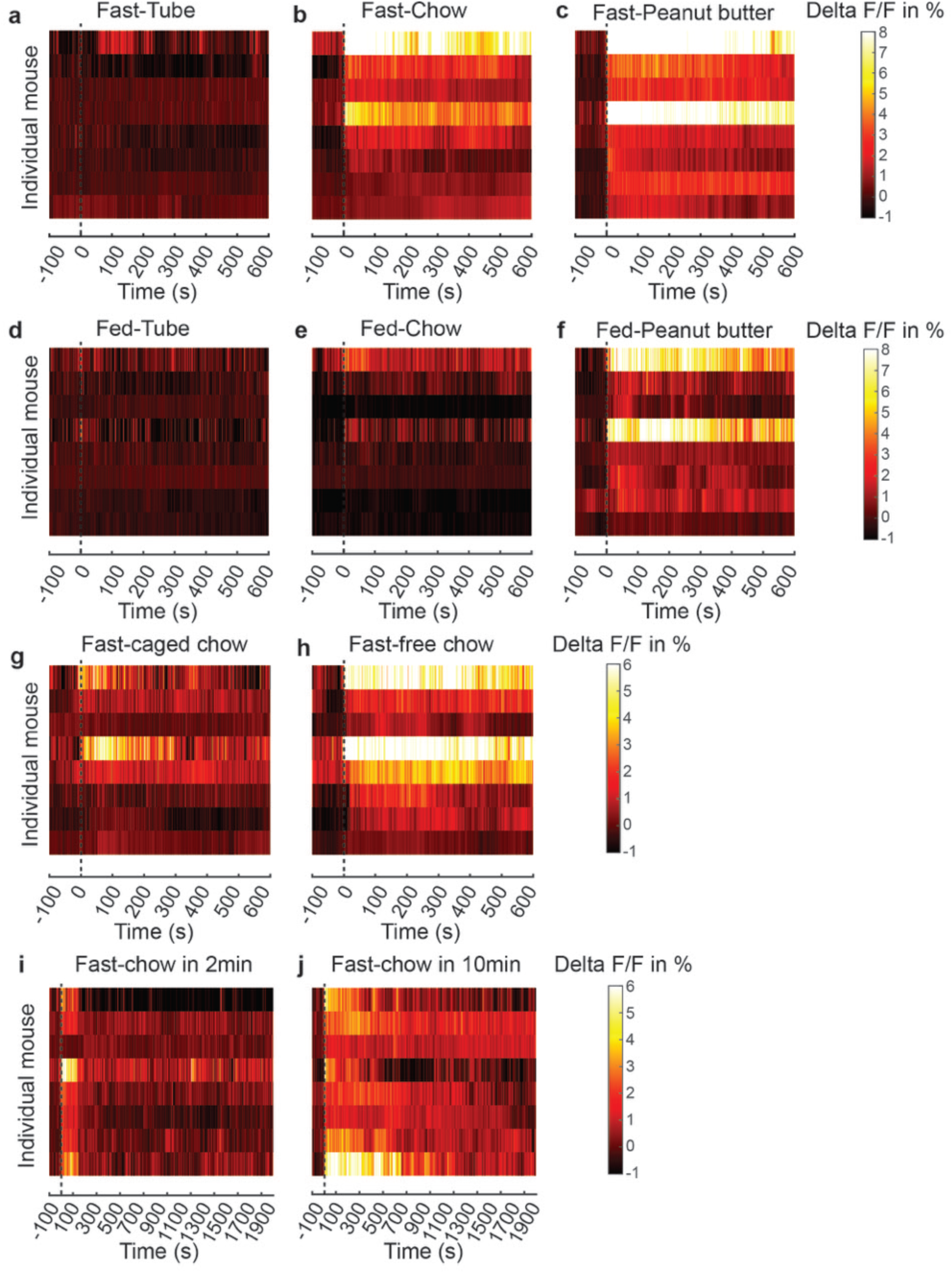
Rapid responses of BNC2 neurons to food cues. **a-c**, Heatmaps of normalized delta F/F in individual overnight fasted mice aligned to the time of the presentation of an inedible tube (**a**), chow (**b**), or peanut butter (**c**). **d-f,** Heatmaps of normalized delta F/F in individual ad-libitum-fed mice aligned to the time of the presentation of an inedible tube (**d**), chow (**e**), or peanut butter (**f**). **g,h,** Heatmaps of normalized delta F/F in individual overnight fasted mice aligned to the time of the presentation of a chow inside the container (**g**) and outside the container (**h**). **i,j,** Heatmaps of normalized delta F/F in individual overnight fasted mice aligned to the time of the presentation of chow for 2 minutes (**i**) and 10 minutes (**j**).

**Extended Data Fig. 5:**
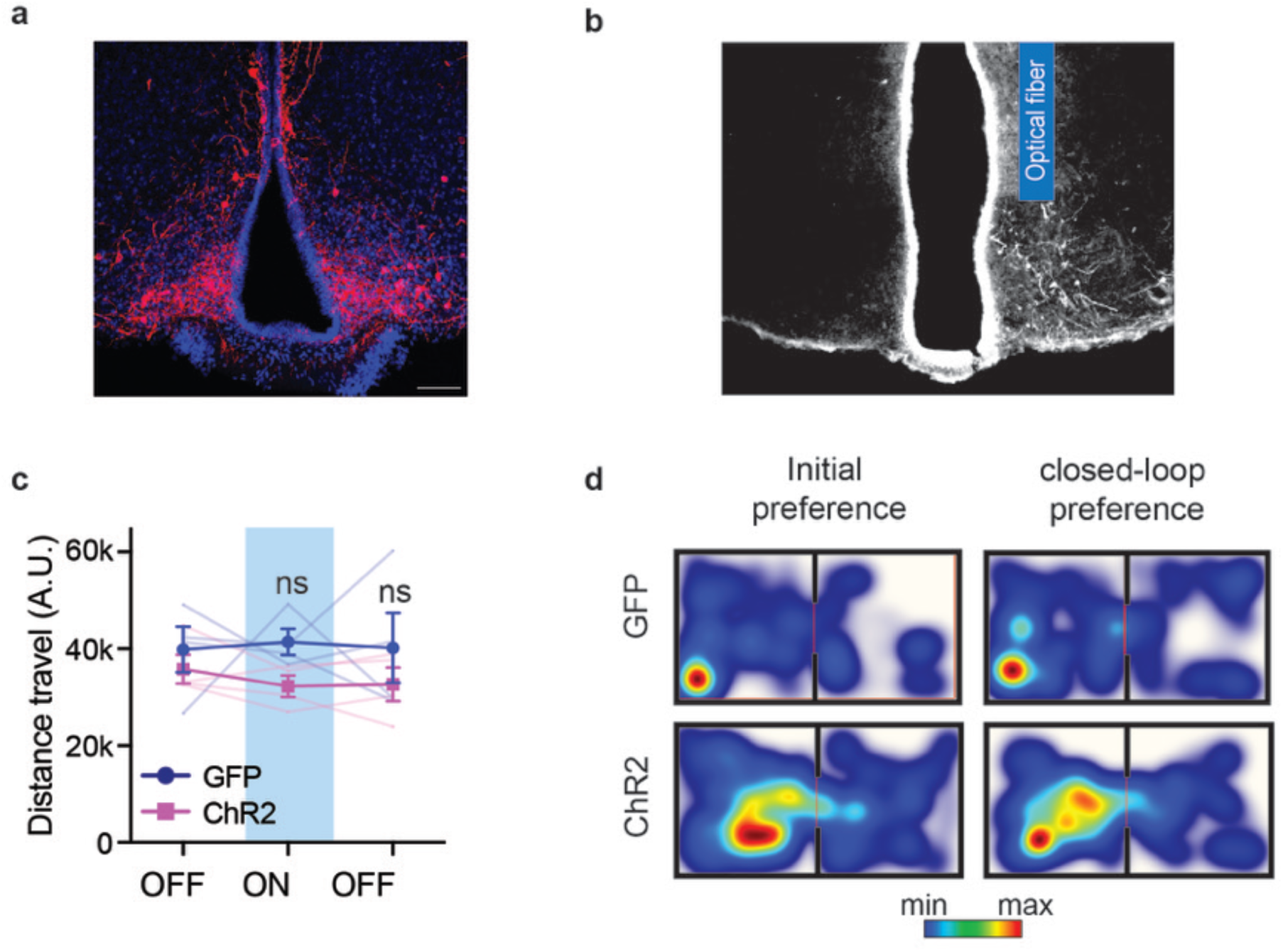
BNC2 neuron activation does not affect locomotion and place preference in sated mice. **a**, Representative image showing bilateral hM3Dq-mcherry expressing (red) in ARC BNC2 neurons. Blue, DAPI. Scale bar, 100 µm. **b,** Representative image showing unilateral ChR2-GFP expression in ARC BNC2 neurons and optical fiber implant site. **c,** Distance travel of mice before, during, and after the activation of laser stimulation in overnight-fasted mice (N = 4 mice per group). **d,** Heatmaps of time spent in each chamber of sated adult male BNC2-Cre mice injected with GFP or ChR2 during the initial phase and the closed-loop paired phase. Data are presented as mean ± SEM. ns, not significant. Details of the statistical analysis are provided as Source Data.

**Extended Data Fig. 6:**
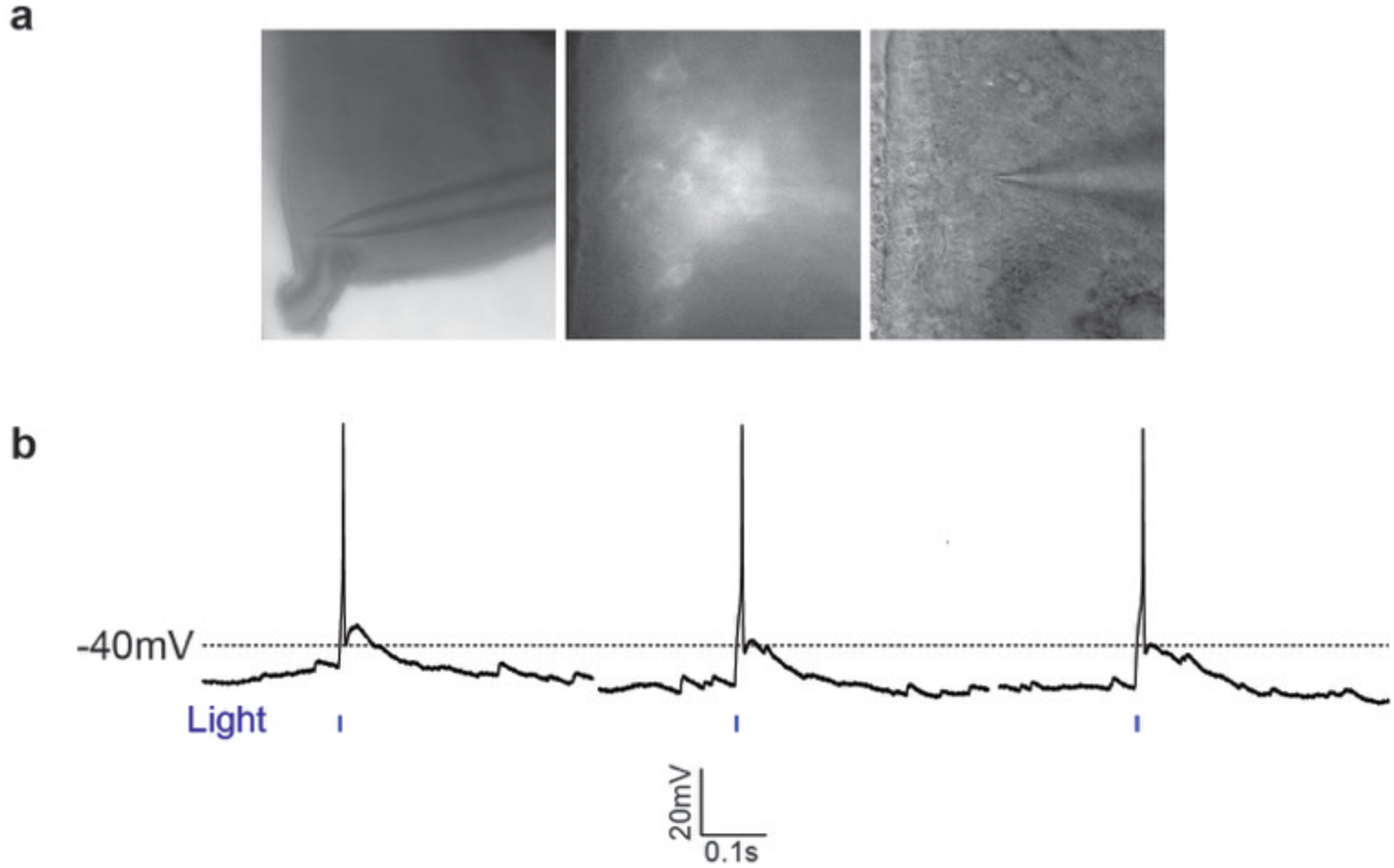
Light-induced activation of NPY neurons. **a**, Representative images of patched ChR2-GFP-labeled NPY neurons. **b,** Detection of light-induced APs in ChR2-GFP-labeled NPY neurons.

**Extended Data Fig. 7:**
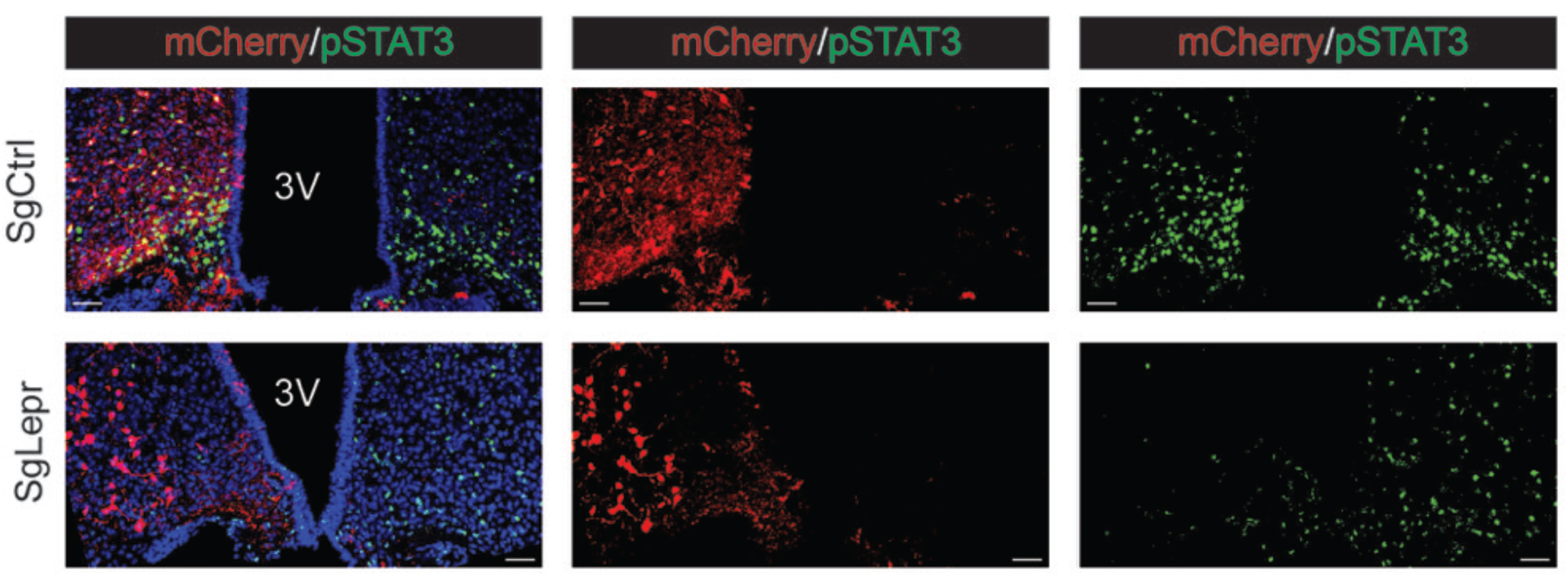
High efficiency of LepR deletion mediated by sglepr. Brain slices from overnight fasted adult male BNC2-Cre::LSL-Cas9-GFP mice, injected with sgCtrl or sglepr into the unilateral ARC, were subjected to mCherry and pSTAT3 immunostaining 3 hours after leptin injection. Scale bar, 50 µm.

**Extended Data Fig. 8:**
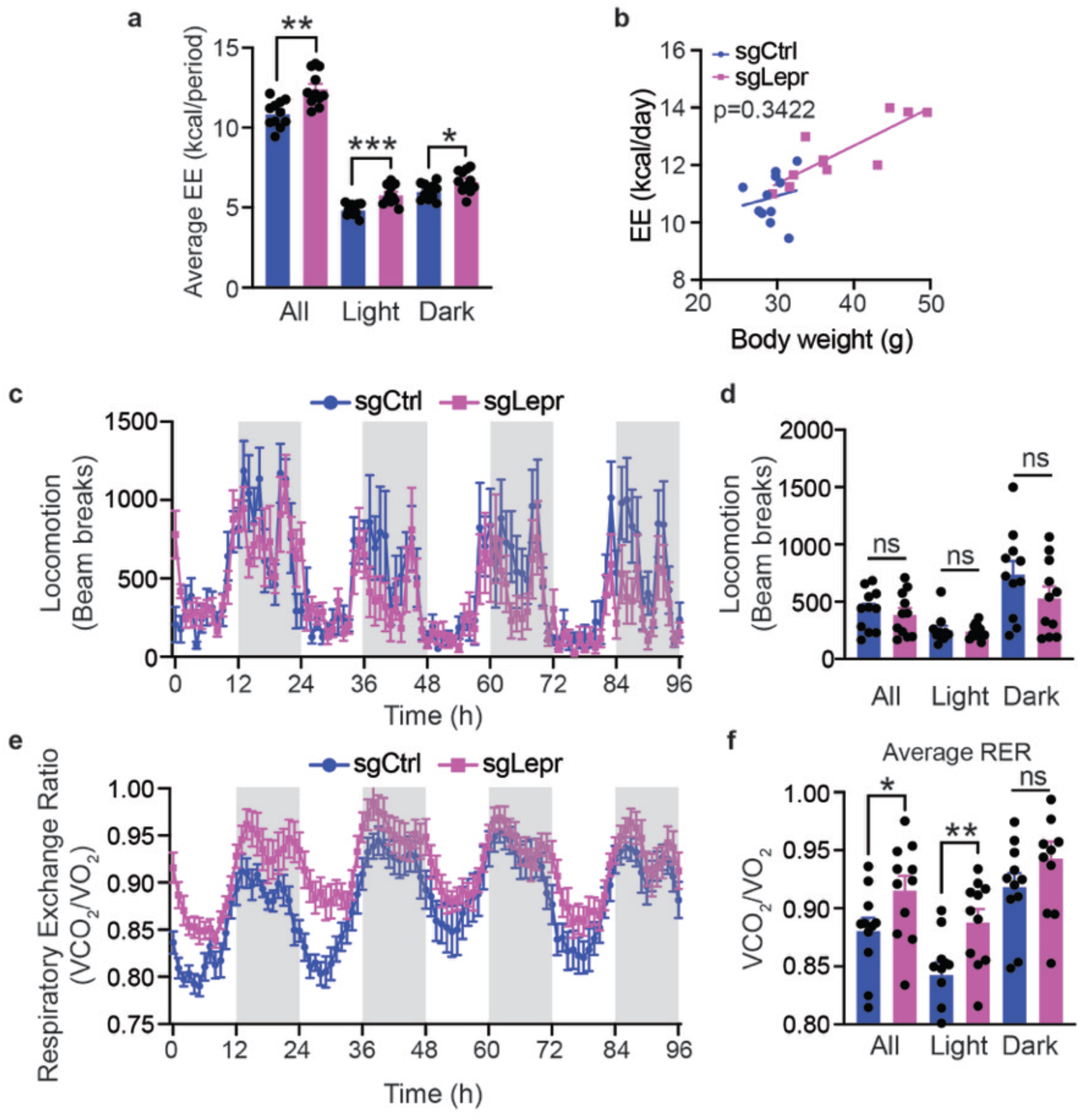
LepR knockout in BNC2 neurons causes higher RER. **a**, Quantification of EE of the two groups (N = 11 mice per group). **b,** Regression analysis of EE vs. total mass from **a** (N = 11 mice per group). **c,** Locomotion of the two groups of mice. **d,** Quantification of locomotion of the two groups (N = 11 mice per group). **e,** Respiratory exchange ratio (RER) of the two groups of mice. **f,** Quantification of average RER of two groups of mice (N = 11 mice per group). Data are presented as mean ± SEM. ns, not significant; *p<0.05. **p<0.01; ***p<0.001. Details of the statistical analysis are provided as Source Data.

**Extended Data Fig. 9:**
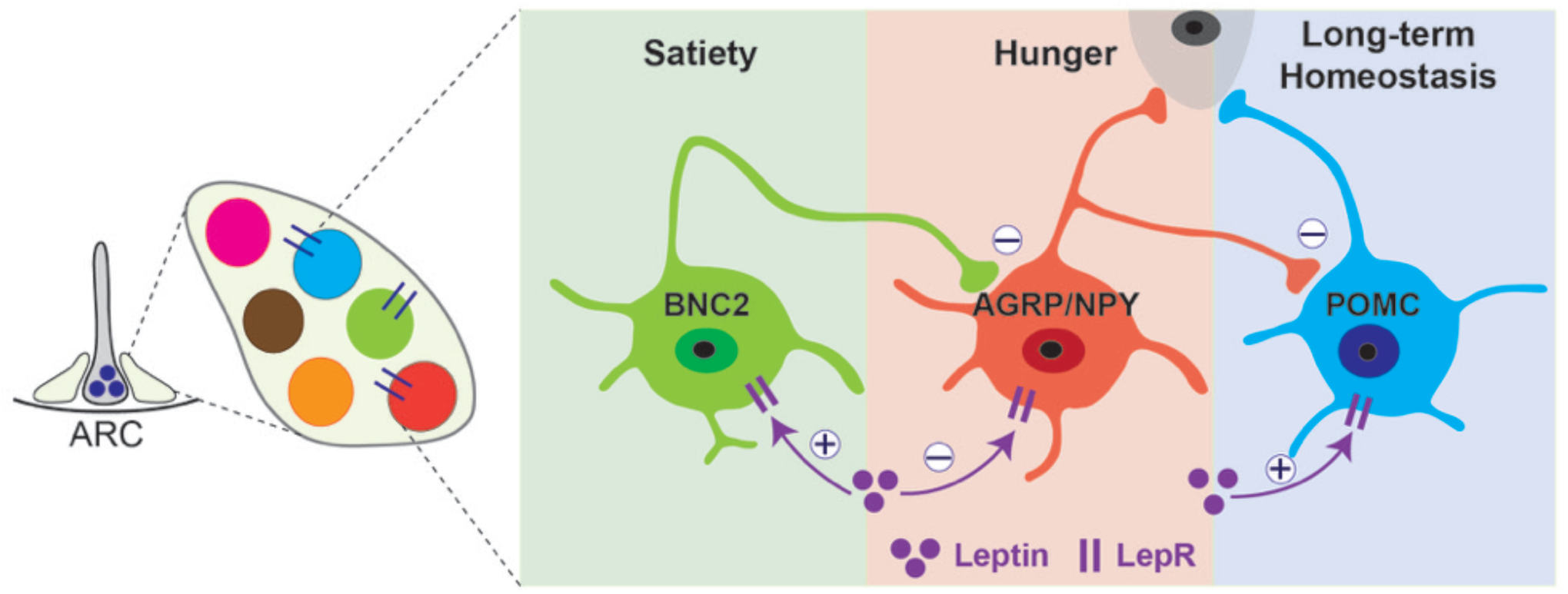
Model of leptin regulation of feeding and energy balance in ARC microcircuits. Leptin-activated BNC2 neurons acutely promote satiety by suppressing AGRP/NPY neurons, which rapidly stimulates hunger. POMC neurons restrict weight gain by modulating energy balance over the long term with minimal short-term effects on food intake.

